# VelA and LaeA are key regulators of the *Epichloë festucae* transcriptomic response during symbiosis with perennial ryegrass

**DOI:** 10.1101/835249

**Authors:** M. Rahnama, P. Maclean, D.J. Fleetwood, R.D. Johnson

## Abstract

VelA (or VeA) is a key global regulator in fungal secondary metabolism and development which we previously showed is required during the symbiotic interaction of *Epichloë festucae* with perennial ryegrass. In this study, comparative transcriptomics analyses of Δ*velA* mutant compared to wild type *E. festucae*, under three different conditions (in culture, infected seedlings and infected mature plants) were performed to investigate the impact VelA on the *E. festucae* transcriptome. These comparative transcriptomics studies showed that VelA regulates the expression of genes encoding proteins involved in membrane transport, fungal cell wall biosynthesis, host cell wall degradation and secondary metabolism, along with a number of small secreted proteins and a large number of proteins with no predictable functions. In addition, these results were compared with previous transcriptomics experiments studying the impact of LaeA, another key global regulator of secondary metabolism and development that we have shown is important for the *E. festucae*- perennial ryegrass interaction. The results showed that although VelA and LaeA regulate a sub-set of *E. festucae* genes in a similar manner, they also regulated many other genes independently of each other suggesting specialised roles.

## Background

*Epichloë* endophytes are symbiotic fungi that systematically colonize intracellular spaces of cool-season grasses of the sub-family Pooideae [1-3]. Most of these symbiotic interactions are mutually beneficial with the plant providing the endophyte with nutrients, and the endophyte protecting host plants from a range of biotic and abiotic stresses [4, 5]. The bioprotective effects of *Epichloë* endophytes are mediated by *in planta* production of four different well characterised classes of alkaloids: indole-diterpenes, ergot alkaloids, lolines and peramine [6].

During this mutualistic interaction, the fungus colonises most of the upper ground parts of the plant in a tightly regulated and synchronized manner [7]. Hyphae only grow in the intercellular spaces, from the meristem to the inflorescence, except for vascular bundles [2, 7]. Also, fungal hyphae continue to grow during host cell growth but cease growth when the host stops growing, although hyphae remain metabolically active [8]. This results in a seldom-branched intercellular network of hyphae, parallel to the leaf axis with hyphae tightly linked with the walls of neighbouring plant cells [7]. A recent study has suggested that this hyphal network is a result of coordinated hyphal growth and death [9].

Although the molecular mechanisms that regulate the mutualistic interaction between *E. festucae* and perennial ryegrass is still largely unknown a number of studies have shown the importance of genes required for hyphal anastomosis (soft gene [10]), fungal biology and development (*velA* gene [11] and *laeA* gene [12]), localised production of reactive oxygen species (required to maintain hyphal polarity [13-16]) and iron homeostasis (*sidN* gene [17, 18]). Although it is not clear how *Epichloë* suppresses or avoids plant responses to establish and maintain a compatible interaction, recent studies have suggested that reducing the chitin inducible host defence response, by altering fungal cell wall chitin, is a possible mechanism [19, 20].

Transcriptomics has also been used to investigate the *Epichloë* interaction with grasses. These studies mostly focused on comparing endophyte free grasses with infected grasses [21-23]. In 2010, the interaction of *Epichloë festucae*/*Lolium perenne* was investigated by comparative transcriptomics of wild type infected plants versus a mutant deleted in the stress-activated MAP kinase gene, Δ*sakA*, - infected plants by Illumina mRNA sequencing. In this study, around 11% of *E. festucae* genes were differentially expressed, with around 75% of these genes up-regulated in Δ*sakA*-infected plants compared with wild type infected plants[24]. In 2015, the same group investigated the *Epichloë festucae*/*Lolium perenne* interaction by using three fungal mutants that cause incompatible interactions in *L. perenne*: Δ*proA*, which encodes a C6-Zn transcription factor that is essential for sexual fruiting body maturation in *Sordaria macrospora*, Δ*noxA*, encoding the NADPH oxidase A, and Δ*sakA*. Three comparative transcriptomics analyses of infected perennial ryegrasse with each of these mutants compared to wild type *E. festucae* infected plants was performed by RNA-seq. 182 genes were found that were differentially expressed in all three comparisons and these were proposed as the core fungal gene set distinguishing mutualistic from antagonistic symbiotic states [20].

The velvet gene, *veA* or *velA*, is a member of the velvet family of genes that normally include four members, *velA, velB, velC* and *vosA*. VelA is an important conserved fungal regulator of a variety of growth and developmental characteristics [25]. In *E. festucae*, we showed that *velA* is required for fungal biology and development and establishing and maintaining the mutualistic interaction of the fungus with its host perennial ryegrass [11].

Although, based on a crystal structure and yeast one-hybrid experiments with the VosA protein, it appears that the velvet domain mediates DNA binding the biochemical mechanism through which VelA exerts control over gene expression is not fully described [26]. Comparative transcriptomics has been used to elucidate the regulatory effects of different genes on fungal transcriptome profiles. Studies of Δ*velA* differential expression compared with wild type fungi in different conditions have mostly used microarrays. These studies showed that VelA regulates genes involved in different fungal developmental and metabolism processes, coordinating with functional analysis of the mutants. These processes include secondary metabolism and morphogenesis in *Penicillium chrysogenum* [27], aflatoxin biosynthetic genes in *Aspergillus flavus* [28] and fungal development and secondary metabolism in *Fusarium fujikuroi* [29]. In *A. fumigatus* and *Aspergillus nidulans* VelA is a global regulator of secondary metabolism [30, 31]. In *A. flavus* it was shown that VelA regulates a broad range of genes, especially secondary metabolism, with 28 of 56 predicted secondary metabolite gene clusters differentially expressed [32]. In *Fusarium graminearum* VelA is involved in regulating various cellular processes [33]. In *Botrytis cinerea* a comparative transcriptomics analysis revealed VelA regulatory effects on fungal genes in a pathogenic interaction with its host plant *Phaseolus vulgaris* including transporters, glycoside hydrolases and proteases [34]. A further study of this fungus growing on solid medium showed that VelA was involved in the regulation of genes encoding SM-related enzymes, carbohydrate-active enzymes, and proteases [35].

LaeA is a global fungal regulator and a predicted interaction partner of VelA which we recently reported is required for *E. festucae* metabolism and development and for establishing and maintaining a successful symbiotic interaction with perennial ryegrass [12]. In addition, a comparative transcriptomics analyses during the early stages of interaction of inoculated perennial ryegrass seedlings with the Δ*laeA* mutant and wild type *E. festucae* suggested a regulatory role for LaeA on the expression of genes for plant cell wall degradation, fungal cell wall composition, secondary metabolism and small-secreted proteins [12, 37].

Based on the knowledge of regulatory roles of VelA on different fungal transcriptome profiles and our findings of VelA importance in *E. festucae* metabolism and development and successful symbiosis [11], we hypothesised that VelA may be involved in regulating the *E. festucae* transcriptome. In this study, a set of comparative transcriptomics analyses of Δ*velA* mutants versus wild type *E. festucae* in culture, in infected seedlings and in infected mature plants was performed. In addition, these results were also compared to the transcriptomics analysis of Δ*laeA* versus wild type *E. festucae* in seedlings [12, 37] and in culture. We identified a specific set of VelA regulated genes that define possible processes required for this symbiotic interaction.

## Results

### Choosing the conditions for transcriptomics studies

The regulatory effects of VelA on *E. festucae* were determined using comparative transcriptomics analysis of wild type and Δ*velA* mutant strains grown under three different conditions; in culture, during the early stage of infection (in seedlings) and in mature infected plants (Table 1). These results were also compared to the transcriptomics analysis of Δ*laeA* versus wild type *E. festucae* in seedlings [12, 37] and in culture.

**Table 1.** Different conditions and comparisons that used for the transcriptomics analysis. WT, wild type; D, dark; L, light; PRG, perennial ryegrass.

| Conditions | In culture (IC) |  | In seedling (S) |  | In planta (IP) |  |
| --- | --- | --- | --- | --- | --- | --- |
| Host | PDA medium covered by cellophane |  | PRG seedlings |  | Mature infected PRG |  |
| Light condition | D |  | 8h L&16h D |  | 8h L&16h D |  |
| Time | 14 d |  | 14 dpi |  | 3 month post inoculation |  |
| *E. festucae* strains and comparisons | WT | $\Delta velA$ | WT | $\Delta velA$ | WT | $\Delta velA$ |

In order to choose the best time for the transcriptomics study, *velA* expression was examined in wild type *E. festucae* under the three different conditions (in culture, in seedlings and in mature infected plants) employed.

As we showed previously, the highest *velA* expression in culture is under nutrient deprivation (water agar plates) and the lowest expression is in nutrient replete medium (PDA) [11]. Similar results were observed for *laeA* expression in different media [12]. Because for artificially infecting ryegrass plants with *Epichloë*, normally fungi that is growing for two weeks in PDA medium use as inoculum, for in culture transcriptomics study also same fungi in PDA medium was used.

For the seedling study, expression of the *velA* gene was examined after inoculating seedlings with wild type *E. festucae*, growing in both the light and dark, at four time points post-inoculation (24 HPI, 2.5 DPI, 6 DPI and 14 DPI). Expression of *velA* increased over time and showed the highest expression 14 DPI. At all-time points, higher expression was detected from seedlings growing under light compared to the dark (Fig1a). As we showed before, the highest *laeA* expression was also at 14 DPI although it was not a gradual increase as was the case with *velA* [12]. In addition to *velA* and *laeA* having the highest expression at 14 DPI, this time point also correlated with when Δ*velA* and Δ*laeA* mutant inoculated seedlings began to die [11, 12]. Because of these reasons, 14 DPI was chosen to study the requirement of *velA* and *laeA* on the expression profile of *E. festucae* during infection of perennial ryegrass.

In mature plants, as we showed before, old blades have the highest *velA* expression compared to other tissues [11] so this was also used in the current transcriptomics study.

The relative expression of *velA* and *laeA* in culture, in seedlings and in mature plants was assessed using qRT-PCR. *velA* showed higher expression in seedlings compared to mature plants. This is opposite to the expression of *laeA* under the same conditions suggesting that *velA* and *laeA* exert different effects on the early stage of infection (Fig. 1b).

**Fig. 1.**
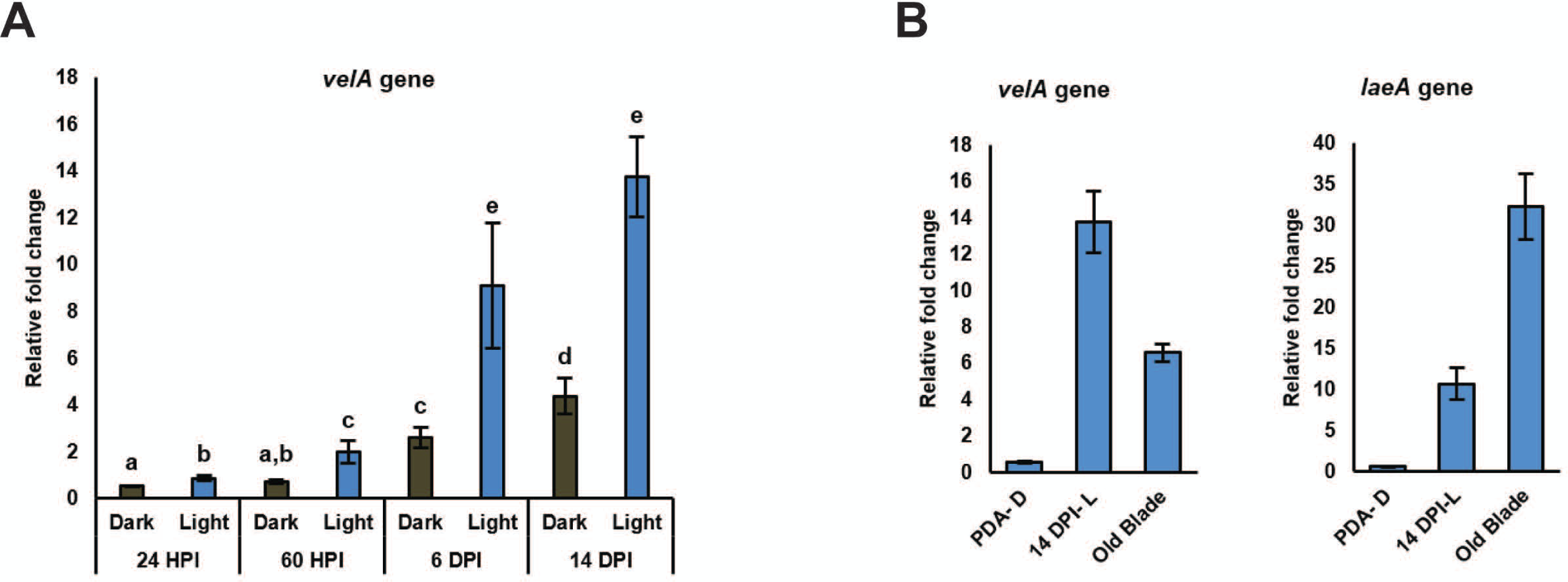
Relative expression of *velA* and *laeA* genes measured by qRT-PCR. a) Relative expression of *velA* in wild type *E. festucae* at different times after inoculating seedlings (HPI: hours post inoculation; DPI: days post inoculation). b) Relative expression of laeA from some chosen conditions in vitro, in culture (PDA- D: potato dextrose agar under dark), seedlings (14 DPI- L: days post inoculation under light) and mature plants (old blade). Expression values were normalised to that of the gamma actin and 60S ribosomal protein L35. Primers used in the analysis are listed in Table S1. Bars represent standard error of the mean calculated from three biological replicates. Results were analysed using the MANOVA Tukey’s test and statistically significant results are indicated with different letters.

### General description of RNA-sequencing results

Higher numbers of reads were generated from in culture samples compared to seedlings and mature plants (Fig. S1a and Table S2). Although similar numbers of reads were generated for both seedling and mature plant samples (Fig S1a) only 1.84% of total reads from mature plants mapped to the *E. festucae* genome compared to 6.28% from seedlings (Table S2) which is indicative of the higher fungal biomass (relative to plant tissue) in seedlings.

Multidimensional scaling (MDS) analysis of the 1,000 most highly expressed genes shows that all in planta replicate samples clustered very close to each other but were separated from in culture samples (Fig. S1b).

Genes with equal or greater than two times fold changes and a FDR (false discovery rate) equal or less than 0.05 were counted as differentially expressed genes (DEGs) (Fig 2a). The percentages of DEGs observed in the S Δ*velA*-WT comparison was higher than other *velA* and *laeA* related comparisons. In S Δ*velA*-WT comparison 5.1 and 1.3 times more DEGs compared to IC Δ*velA*-WT and IP Δ*velA*-WT comparisons were observed which indicates a significant impact of the growth conditions on the number of DEGs in the Δ*velA* mutant. Only 10% and 23.5% of DEGs (29 and 69 genes) in the S Δ*velA*-WT comparison was common with DEGs in the IC Δ*velA*-WT and IP Δ*velA*-WT comparisons, respectively (Fig 2b).

**Fig. 2.**
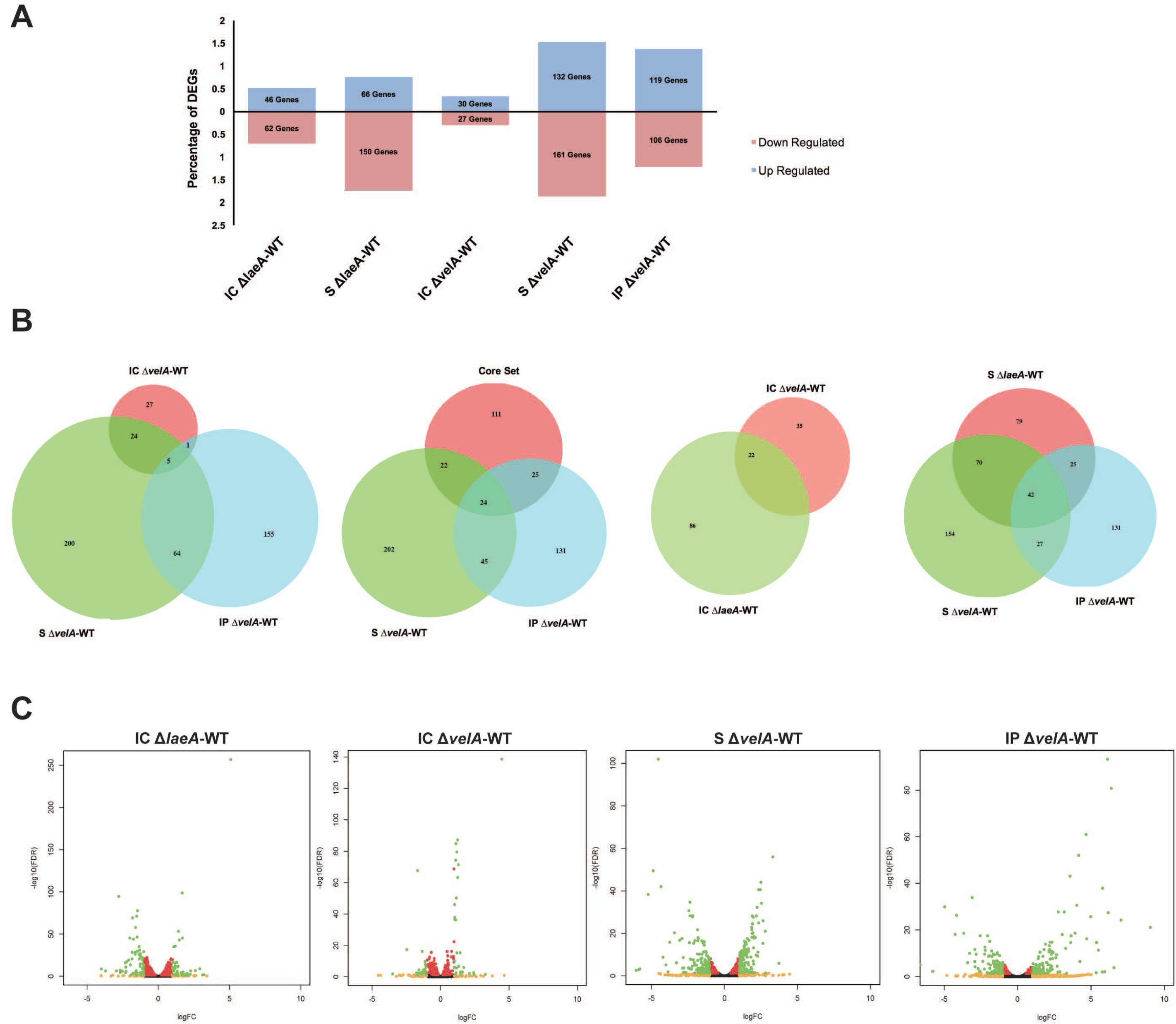
Percentage and distribution of differentially expressed genes in different comparisons and different conditions. a) The bar chart shows the percentage of DEGs that up or down-regulated in different comparisons. b) Venn diagrams show the common DEGs genes between different comparisons. c) Volcano plots show distribution of log_2_ of fold changes (logFC) and significantly (–log_10_ of FDR) in different comparisons. Black dots: FDR>0.05, Red dots: FDR<0.05, orange dots: logFC>1, green dots: FDR<0.05 & logFC>1.

In the IC Δ*laeA*-WT comparison there was almost 2 times more DEGs compared to IC Δ*velA*-WT and only 15.38% (22 genes) of them are common (Fig. 2a and b). DEGs in S Δ*laeA*-WT are around 26% less than S Δ*velA*-WT but similar to IP Δ*velA*-WT (Fig. 2a). More than 69% of the DEGs in S Δ*laeA*-WT are downregulated but in both S Δ*velA*-WT and IP Δ*velA*-WT a similar percentage of DEGs are up and down-regulated (Fig. 2a). DEGs in S Δ*laeA*-WT comparison had 52% and 31% (112 and 67) in common with S Δ*velA*-WT and IP Δ*velA*-WT comparisons, respectively, which indicates different regulatory effects of LaeA and VelA on the fungal transcriptome during interaction with its host (Fig. 2b).

There are 182 DEGs common in transcriptomes of three *E. festucae* mutants (Δ*proA*, Δ*noxA*, and Δ*sakA*) which were proposed to form a core set of *Epichloë* genes that distinguished mutualistic from antagonistic symbiotic states [20]. Our results for the S Δ*velA*-WT and IP Δ*velA*-WT comparisons showed that only 46 and 49 genes were in common with the proposed core set from Eaton *et al*, respectively (Fig. 2b). Thus, not all the core genes identified in the Eaton *et al.* (2015) study are required for distinguishing mutualistic from antagonistic symbiotic states.

IP Δ*velA*-WT fold change range (524.7 to −99.8) was 13 and 7 times greater compared to IC Δ*velA*-WT (22.3 to −11.4) and S Δ*velA*-WT (13.2 to −66.2) comparisons, respectively (Fig. 2c and Fig. S1c). Interestingly, IC Δ*laeA*-WT (33.7 to −16.0) and S Δ*laeA*-WT (12.6 to −56.3) showed similar fold change ranges with IC Δ*velA*-WT and S Δ*velA*-WT, respectively (Fig. 2c and Fig. S1c).

### Gene Ontology (GO) enrichment analysis on DEGs

DEGs were classified based on their primary functions into ‘Molecular Function’ and ‘Biological Process’ GeneOntology (GO) categories (Fig. S2 and Table S3). 55-72% of DEGs in different comparisons were not aligned with any GO category so it was decided that GO was not a good method to study the functions of DEGs in this study.

### Functional annotations of differentially expressed *E. festucae* genes

To further investigate the DEGs functions, BLAST analyses were performed against different databases including Uniprot, Swissprot, InterProScan and KEGG and their putative functions were manually determined. In 36.8% (21), 46.1% (135), and 46.2% (104) of DEGs in IC Δ*velA*-WT, S Δ*velA*-WT, and IP Δ*velA*-WT comparisons, respectively, no significant BLAST hits (p-value ≤ E-20) were found. This suggests that many of the genes regulated by VelA are likely to be unique to *E. festucae*. Similar numbers of DEGs genes were found with no significant hits in IC Δ*laeA*-WT and S Δ*laeA*-WT with 35.2% (38) and 22.2% (48), respectively.

### Changes in expression of genes encoding orthologues of velvet family members

We previously reported orthologues of velvet family members in *E. festucae*: VelA (EfM3.049680), VelB (EfM3.023360), VelC (EfM3.009960), VosA or VelD (EfM3.010530) and LaeA (EfM3.069170) [11, 12]. Differential expression of these genes in different comparisons showed that *laeA* expression in IC Δ*velA*-WT and S Δ*velA*-WT comparisons was up regulated. However, *velA* expression did not change significantly in Δ*laeA* mutant related comparisons (Table S4) which suggests that whilst VelA negatively regulates the expression of the *laeA* gene, LaeA does not have any significant influence on the expression of *velA*. The expression of other members of the velvet family did not changed significantly in any of the Δ*velA* or Δ*laeA* mutants (Table S4).

### DEGs in different functional categories

Further analysis focussed on five areas of fungal cellular function and metabolism: fungal cell membrane transporters, fungal cell wall biosynthesis, host cell wall degradation, secondary metabolism and small-secreted proteins. These five areas include most of the DEGs that were previously reported as important in the *E. festucae*/*L. perenne* interaction [20].

#### 1. Changes in expression of genes encoding membrane transporters

DEGs that encode transporters are summarised in table 2. For the in culture comparison, there were a small number of DEGs (4) with transporter activity but it significantly increased (11-15) in S Δ*velA*-WT, S Δ*laeA*-WT and IP Δ*velA*-WT comparisons. This indicates a greater regulatory effect of VelA and LaeA on the expression of *E. festucae* genes encoding membrane transporters during fungal interaction with ryegrass. Higher numbers of DEGs with transporter activity was observed in both *velA* related comparisons (13 and 15 for S Δ*velA*-WT and IP Δ*velA*-WT comparisons) with greater fold changes (2.1-12.5) than S Δ*laeA*-WT comparison (11 DEGs with 2 - 4.8 fold change range). This suggests stronger regulatory effects of VelA compared to LaeA on the expression of *E. festucae* membrane transporters (Table 2).

**Table 2.**
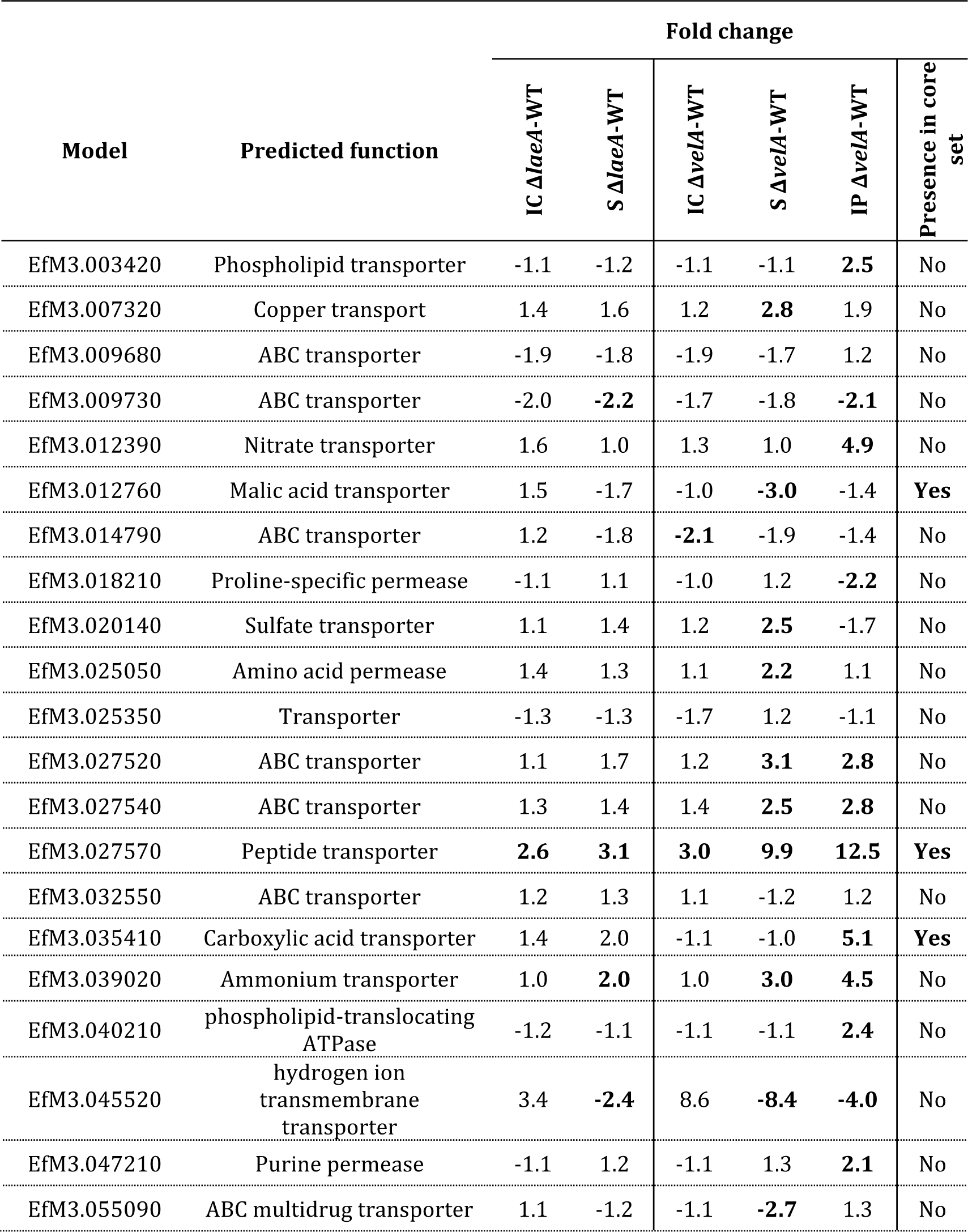
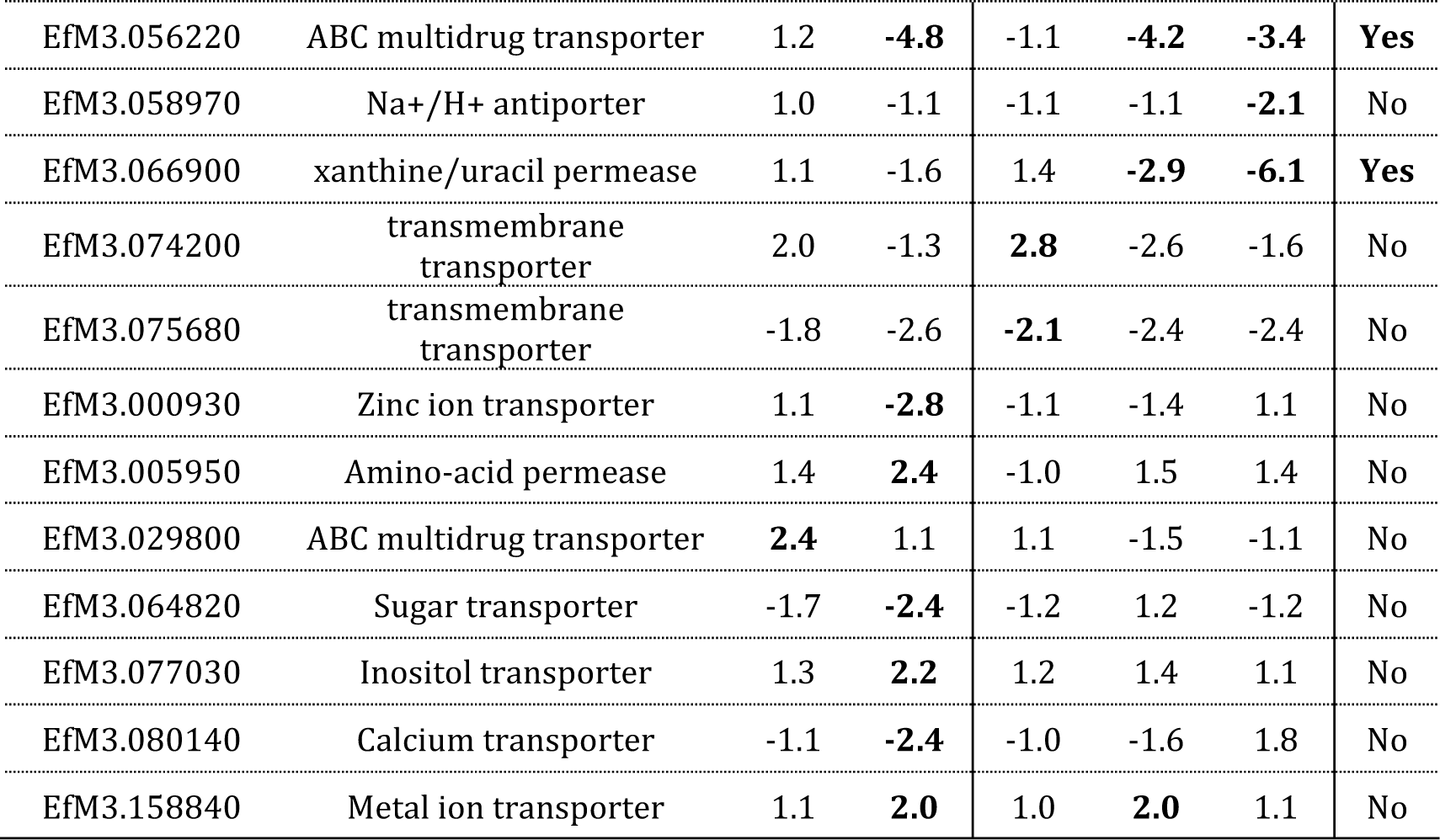
DEGs with transporter activity in all different comparisons

The genes with the highest fold changes (more than four times) were involved in transporting different compounds such as nitrogen, peptides, hydrogen ion and carboxylic acid. There are two genes, EfM3.012390 and EfM3.039020, involved in nitrogen transport. EfM3.012390, which was only differentially expressed in the IP Δ*velA*-WT comparison (4.9-fold up-regulated), is a homologue of a nitrate transporter, CrnA, from *Emericella nidulans* and has been shown to be involved in nitrogen metabolite repression [38]. EfM3.039020, which was up-regulated in both S Δ*velA*-WT and IP Δ*velA*-WT comparisons is a homologue of an ammonium transporter, Amt1, from *Schizosaccharomyces pombe* [39]. EfM3.027570 was up-regulated in both S Δ*velA*-WT and IP Δ*velA*-WT comparisons and is a homologue of a peptide transporter, PTR2, from *Stagonospora nodorum* [40].

These results suggest that Δ*velA* and Δ*laeA* mutants may be nutrient starved during interaction with seedlings or mature plants.

#### 2. Changes in expression of genes encoding enzymes with host cell wall degrading activity

Fundamentally, plant cell walls are made of embedded cellulose microfibrils in a matrix of pectin, hemicellulose and cell wall associated proteins [41]. To look specifically at the enzymes with plant cell wall degrading characteristics, the Fungal PCWDE (Plant Cell Wall-degrading Enzymes) Database [42], was utilised. In this database there are 22 known fungal gene families which degrade the plant cell wall [42]. Searching against this database showed that 25 genes in 13 PCWDE families are present in *E. festucae* (Table S5).

In addition, the larger Carbohydrate-Active enZYmes (CAZyme) database [43], which includes the families of enzymes that assemble, modify, or breakdown oligo- and polysaccharides was checked [43]. The presence of 310 genes in *E. festucae* that are homologues with CAZyme families [20] were analysed for DEGs across the different comparisons (Fig. S3). The number of DEGs with homologues to CAZyme families was almost ten times and two times increased in S Δ*velA*-WT compared to IP Δ*velA*-WT and IC Δ*velA*-WT, respectively, and two times increased in S Δ*laeA*-WT compared to IC Δ*laeA*-WT (Fig. S3). These results indicate the importance of VelA and LaeA on the expression of CAZyme genes during the *E. festucae* interaction with its host plant.

All DEGs with plant cell wall degrading activity are summarised in table 3. No DEGs belonging to cellulases, xylanases, cutinases or pectinases were detected for the in-culture comparisons. No DEGs with cellulose degradation activity was detected for S Δ*velA*-WT and IP Δ*velA*-WT comparisons but two DEGs with cellulose activity were detected in the S Δ*laeA*-WT.

**Table 3.** DEGs with host cell wall degradation activity. Fold changes showed in bold are statistically significant (FDR≤0.05).

| Model | CAZyme class | Predicted function | Target cell wall component | Fold change |  |  |  |  |
| --- | --- | --- | --- | --- | --- | --- | --- | --- |
| | | | | IC ΔlaeA-WT | S ΔlaeA-WT | IC ΔvelA-WT | S ΔvelA-WT | IP ΔvelA-WT |
| EfM3.049570 | GH13 | Alpha- glucosidase | Cellulose | 1.0 | **-2.1** | -1.2 | -1.5 | -1.2 |
| EfM3.053990 | GH5 | Endoglucanase | Cellulose | 1.5 | **2.1** | -1.1 | 1.1 | 1.3 |
| EfM3.005420 | N/A | Exo-1,4-beta-xylosidase | Hemicellulose | 1.1 | **1.9** | -1.0 | **2.3** | 1.5 |
| EfM3.040190 | GH10 | Endo-1,4-beta-xylanase | Hemicellulose | 1.7 | **3.3** | 1.1 | **3.9** | **83.5** |
| EfM3.037040 | GH62 | Glycosyl hydrolase | Hemicellulose | 1.1 | **4.3** | 1.0 | **4.3** | **69.6** |
| EfM3.008730 | CE8 | Pectin methylesterase | Pectin | -1.3 | 1.7 | 1.0 | **2.0** | 1.4 |
| EfM3.008610 | CE5 | Cutinase | Cutin | **2.5** | 2.9 | 1.2 | -2.0 | 2.9 |
| EfM3.005300 | N/A | Cuticle-degrading protease | Cuticle | 1.0 | **-2.8** | 1.0 | **-5.1** | **-4.1** |

Three DEGs, EfM3.040190, EfM3.005420 and EfM3.037040, with hemicellulose activity were detected (Table 3). Two of them, EfM3.037040 and EfM3.040190, were up-regulated in all S Δ*laeA*-WT, S Δ*velA*-WT and IP Δ*velA*-WT comparisons, with the IP Δ*velA*-WT comparison showing the highest fold change of all DEGs with plant cell wall degradation activity (Table 3). EfM3.037040 is a homologue of xylanase C (XynC) from *Cellvibrio japonicus* which hydrolyses hemicellulose (Kellett et al., 1990). This gene was 4.3, 4.3 and 69.6 fold up-regulated in Δ*laeA*-WT, S Δ*velA*-WT and IP Δ*velA*-WT comparisons, respectively. EfM3.040190 is a homologue of endo-1,4-beta-xylanase 2 (*xyl2*) from *Claviceps purpurea* [44] and was 3.3, 3.9 and 83.5 fold up-regulated in S Δ*laeA*-WT, S Δ*velA*-WT and IP Δ*velA*-WT comparisons, respectively. In addition to these two genes another DEG was observed with hemicellulose activity, EfM3.005420, that is a homologue of exo-1,4-beta-xylosidase (*bxlB*) from *Aspergillus flavus* and which was 2.3-fold up-regulated in S Δ*velA*-WT.

One DEG, EfM3.008730, 2-fold was up-regulated in the S Δ*velA*-WT comparison and encodes a pectinase with homology to pectin methyl esterase (*pme1*) in *Aspergillus aculeatus*. It has been shown that Pme1 in *A. aculeatus* degrades the host plants cell wall [45].

A down-regulated DEG, EfM3.005300, identified from both seedlings and in planta comparison which is a homologue of the cuticle degrading protease from the insect pathogen *Metarhizium anisopliae* [46].

Based on these data it appears VelA regulates *E. festucae* genes that encode proteins with a range of plant cell wall degrading functions during the fungal interaction with its host. Of the surviving plants infected with the Δ*velA* mutant, there was significantly greater hemicellulose degradation activity which may contribute to the observed phenotypes of mutant infected plants [11].

#### 3. Changes in expression of genes encoding proteins involved in fungal cell wall composition

Chitin, glycoproteins and glucan are the main components of the fungal cell wall and these can function as elicitors of plant defence responses [47, 48]. Fungi use a range of enzymes to break down, synthesise or remodel their cell wall [49]. All detected DEGs that encode proteins related to the fungal cell wall are summarised in table 4.

**Table 4.**
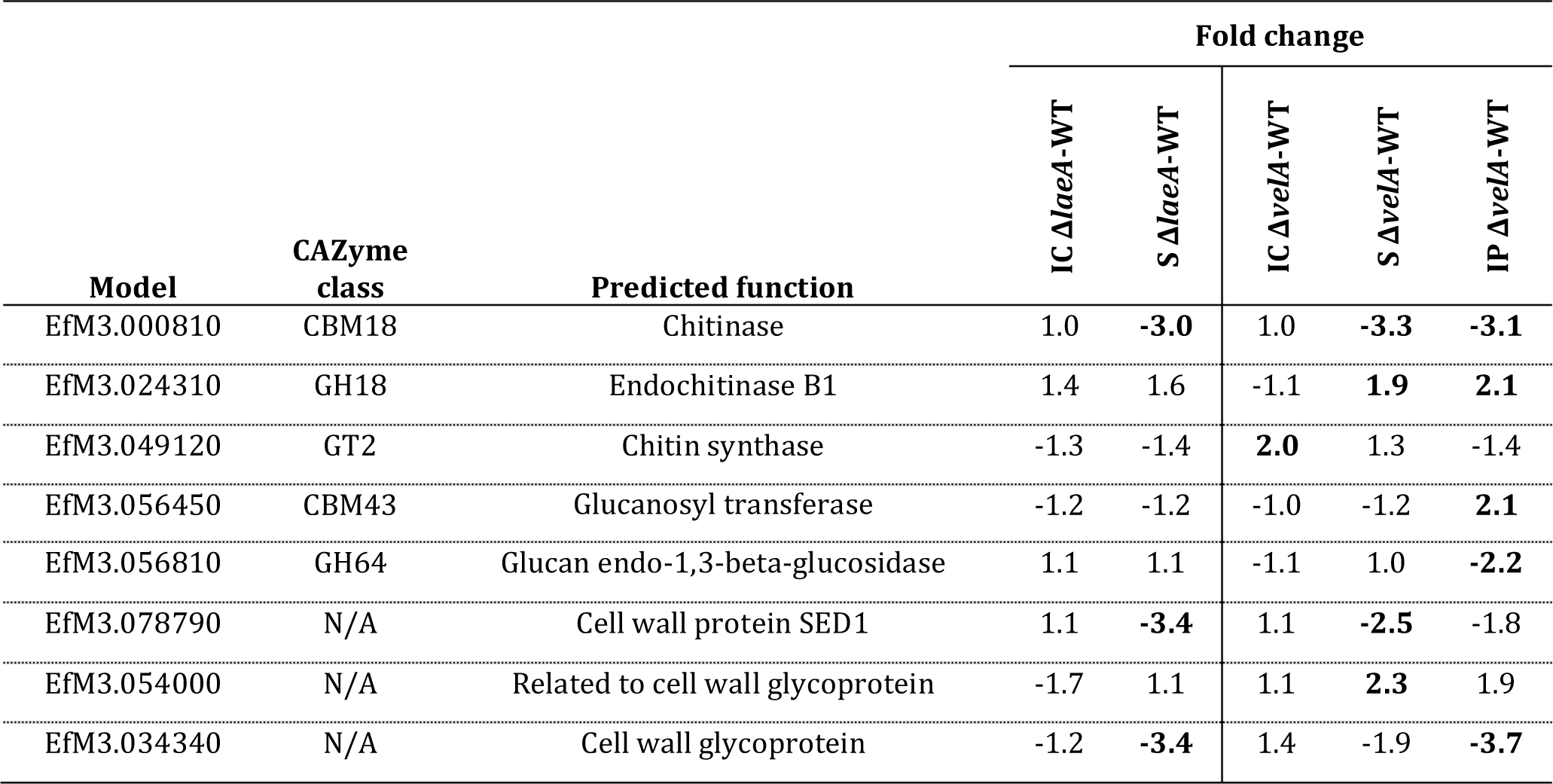
Differentially expressed genes engage in fungal cell wall composition. Fold changes showed in bold are statistically significant (FDR≤0.05).

Enzymes for the synthesis and breakdown of chitin are chitin synthases and chitinases, respectively [49]. EfM3.000810 is a homologue of a chitinase (BDCG_06828) from *Blastomyces dermatitidis* and was 3, 3.3 and 3.1 fold down-regulated in S Δ*laeA*-WT, S Δ*velA*-WT and IP Δ*velA*-WT comparisons, respectively. EfM3.024310 is a homologue of Endochitinase B (*chiB1*) from *Neosartorya fumigata* [50] and was 2.1-fold up-regulated in the IP Δ*velA*-WT comparison. EfM3.049120 is a homologue of chitin synthase 1 (*chs-1*) from *Neurospora crassa* and was 2-fold up-regulated in the IC Δ*velA*-WT comparison. This gene has been shown to play a major role in cell wall biogenesis in this fungus [51].

No DEGs that encode β-1,3-glucan synthase enzymes, responsible for synthesis of fungal cell wall glucan, were found in *E. festucae* but there were two genes, EfM3.056450 and EfM3.056810 that are possibly engaged in breaking down glucan. EfM3.056450 is a homologue of EPD1 from the dimorphic yeast *Candida maltosa*. In this fungus, it was shown that EPD1 is necessary for fungal transition to the pseudohyphal growth form. Deleting this gene leads to a reduction of both alkali-soluble and alkali-insoluble β-glucan levels [52]. EfM3.056450 also has a glucanosyl transferase domain that in some cases has been shown to remodel chains of β −1,3-glucan in the fungal cell wall [53]. This gene is up-regulated 2.1-fold in the IP Δ*velA*-WT comparison. EfM3.056810 is a homologue of glucan endo-1,3-beta-glucosidase from *Cellulosimicrobium cellulans* and was 2.1-fold down-regulated in the IP Δ*velA*-WT comparison. This enzyme was shown to be involved in breaking down fungal cell walls by hydrolysis of cell wall β-1,3-glucosidic linkages [54].

Glycoproteins are another constituent of the fungal cell wall that are made of modified proteins by *N*- and *O*-linked carbohydrates that in lots of examples are glycosyl phosphatidyl inositol (GPI) anchors [49]. There are three genes, EfM3.078790, EfM3.054000 and EfM3.034340, with possible activity towards glycoproteins (Table 4). EfM3.078790 is a homologue of a cell wall glycoprotein *sed1* from *Saccharomyces cerevisiae* that was differentially expressed in seedling comparisons (2.5 fold down-regulated in S Δ*velA*-WT comparison). In *S. cerevisiae* Sed1 is the most abundant cell wall-associated protein in the stationary phase and is necessary for fungal resistance to lytic enzymes [55]. EfM3.054000 is a homologue of a putative cell wall glycoprotein (CPUR_07530) from *Claviceps purpurea* that was 2.3-fold up-regulated in S Δ*velA*-WT comparison. EfM3.034340 is a homologue of the collagen α-2(IV) chain in the nematode *Ascaris suum* and was 3.4 and 3.7-fold down-regulated in the S Δ*laeA*-WT and IP Δ*velA*-WT comparison. Although collagen is best understood in animals it is also detected in fungal fimbriae, long (1-20 µm) and narrow (7 nm) flexuous appendages on the fungal surface, which affect cellular functions such as mating and pathogenesis [56]. This gene is also a homologue of a putative cell wall protein (XP_001270917) in *Aspergillus clavatus*.

One of the fungal phenotypes observed in mature plants infected with the mutant Δ*velA* was formation of intrahyphal hyphae [11]. It has been shown that triggering activation of woronin bodies, important organelles that block the septal pore in response to wounding, increases the formation of intrahyphal hyphae [57]. EfM3.049350 is a homologue of hex protein, a major woronin body protein in *Neurospora crassa* which was up-regulated 2.9 times in the IP Δv*elA*-WT comparison.

These results demonstrate that VelA is a regulator of *E. festucae* cell wall composition during the fungal interaction with ryegrass.

#### 4. *velA* is required for secondary metabolite gene expression and production

In order to investigate the regulatory effects of VelA on secondary metabolism in *E. festucae*, differential expression of genes involved in known alkaloid gene clusters (ergot alkaloids, indole-diterpenes and peramine) were examined across all comparisons. In the in culture comparison, the *lpsB* gene from the ergovaline pathway was up regulated (more than 9 folds) in both Δ*velA* and Δ*laeA* mutants. Another gene from this pathway, *easA*, was also up regulated (2 fold) in Δ*laeA* mutant (Table S6).

Of the 11 genes involved in ergovaline production (*EAS* cluster genes) [58] only 1 gene, *lpsB*, was up-regulated in the IP Δ*velA*-WT comparisons whereas all genes were down-regulated in the S Δ*velA*-WT comparisons, which was similar to the S Δ*laeA*-WT comparison (Table S6, Fig 3a).

**Fig. 3.**
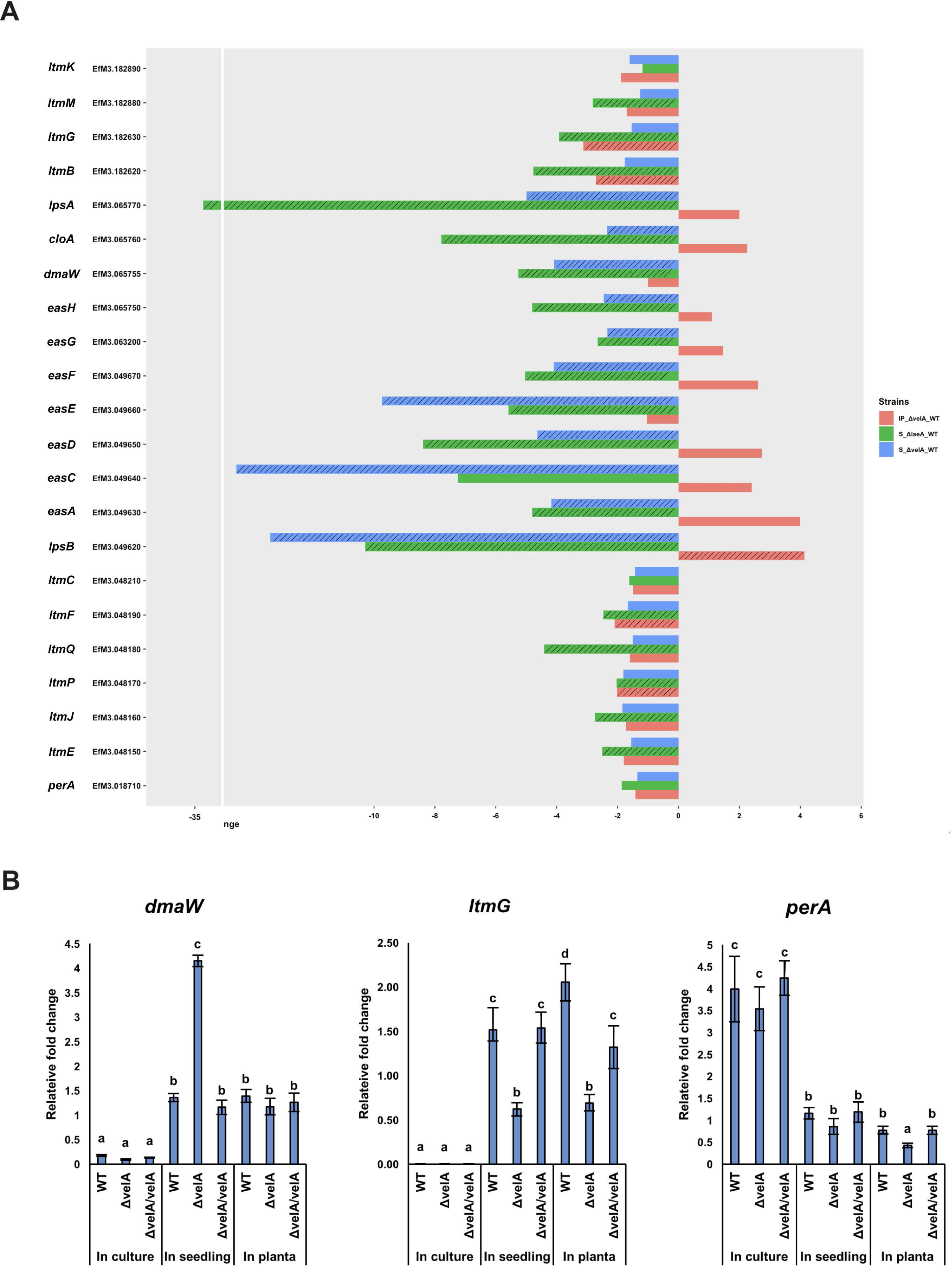
Expression changes of *E. festucae* alkaloid biosynthesis genes in different conditions. a) Expression changes of *E. festucae* alkaloid biosynthesis genes in different comparisons including S Δ*laeA*-WT, S Δ*velA*-WT, and IP Δ*velA*-WT. Dashed bars: fold changes that are statistically significant (FDR≤0.05), non-dashed bars: fold changes that are statistically non-significant (FDR≥0.05). b) Relative expression levels of three representative genes of ergovaline, lolitrem B and peramine biosynthetic genes in wild type, Δ*velA* and Δ*velA/velA* in three different conditions. Results were analysed using the MANOVA Tukey’s and the independent sample T-test and statistically significant results are indicated with same letter are not significantly different.

Of the ten known genes for indole-diterpene biosynthesis, four genes were significantly down-regulated in the IP Δ*velA*-WT comparison. In the S Δ*velA*-WT comparison none of the genes were differentially expressed, which is in stark contrast to the S Δ*laeA*-WT comparison where eight genes were down-regulated. This indicates very different regulatory roles of *velA* and *laeA* genes on fungal secondary metabolite regulation in *E. festucae* (Table S6, Fig. 3a).

The expression of *perA*, the sole gene required for peramine biosynthesis, was not differentially expressed in any comparison (Table S6, Fig. 3a).

The transcriptomics results were validated by qRT-PCR of *dmaW, ltmG* and *perA*, the first gene from the ergovaline, lolitrem B, and peramine pathways, respectively, in all three different conditions using wild type, mutant and complemented strains (Fig. 3b). Results validated the transcriptomics results (Fig. 3b).

To correlate alkaloid pathway gene expression results, different alkaloids concentrations were also measured in infected three-months-old ryegrass plants infected with wild type and Δ*velA* mutant *E. festucae* (Fig. 4a). Of the seven ergot alkaloid compounds that were measured two of them, lysergic acid and elymoclavine, were not detected. For the remaining five compounds, the mean concentrations in the wild type-infected plants were higher than Δ*velA*-mutant infected plants but only ergine was statistically significant (Fig. 4a). Of the 23 indole-diterpene alkaloid compounds that were measured, the mean concentrations of the Δ*velA*-mutant infected plants were higher with 17 of them being statistically significant (Fig. 4a). For peramine the mean concentration in Δ*velA* mutant infected plants was also higher but it was not statistically significant (Fig. 4a).

**Fig. 4.**
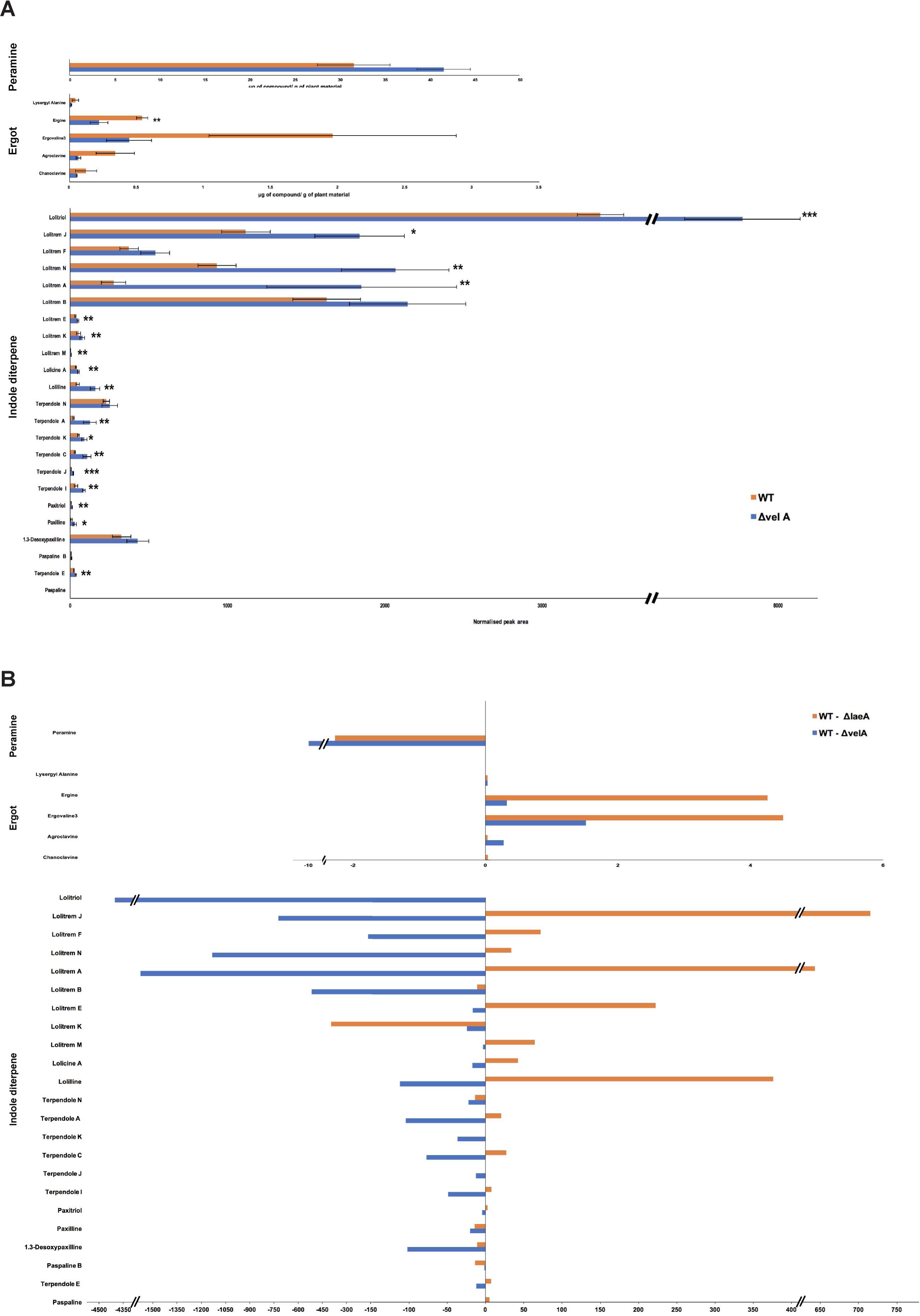
VelA regulates alkaloid production of *E. festucae* in ryegrasses plants. a) Alkaloids’ concentration in three months old ryegrasses infected with wild type and Δ*velA* mutant *E. festucae*. Results were analysed using the independent sample T-test and statistically significant results are indicated (***p<0.001, ** 0.05>p>0.001, * 0.1>p>0.05). b) The differences of mean concertation of the wild type infected plants to the Δ*velA* mutant infected plants and the mean concertation differences of wild type infected plants to Δ*laeA* mutant infected plants.

We previously reported alkaloid production in three-month-old ryegrass plants infected with wild type and Δ*laeA* mutant *E. festucae* [12] (Fig. S7). The differences of mean concertation of wild type infected plants to Δ*velA* mutant infected plants were compared to the mean concentration differences of wild type infected plants to Δ*laeA* mutant infected plants (Fig. 4b). Results show lower concentrations of most of indole-diterpene alkaloids in the Δ*laeA* mutant infected plants which is opposite to that seen for Δ*velA* mutant infected plants (Fig. 4b).

In addition to these 3 alkaloid clusters, 29 other clusters, 191 genes, which putatively encoding secondary metabolites in *E. festucae* Fl1 [6] also were examined (Table S6). Heat maps generated for all 32 clusters in *E. festucae* showed that most of them are down regulated in Δ*velA* mutant infected plants (Fig. 5a), showing that VelA positively regulates secondary metabolite gene expression in *E. festucae*. Of the 32 clusters 12 contained at least one gene differentially expressed in one of the Δ*velA* related comparisons, nine in IP Δ*velA*-WT, ten in S Δ*velA*-WT and three in IC Δ*velA*-WT have DEGs (Fig. 5b, left panel). In the Δ*laeA* related comparisons, also 12 clusters contained at least one gene differentially expressed in one of the comparison, eleven in S Δ*laeA*-WT and five in in IC Δ*laeA*-WT (Fig. 5b, left panel), but these clusters are not all identical to the ones in the Δ*velA* related comparisons. Clusters 20, 29 and 38 are unique to Δ*velA* related comparisons and clusters 10, 35 and 50 are unique to Δ*laeA* related comparisons (Fig. 5b).

**Fig. 5.**
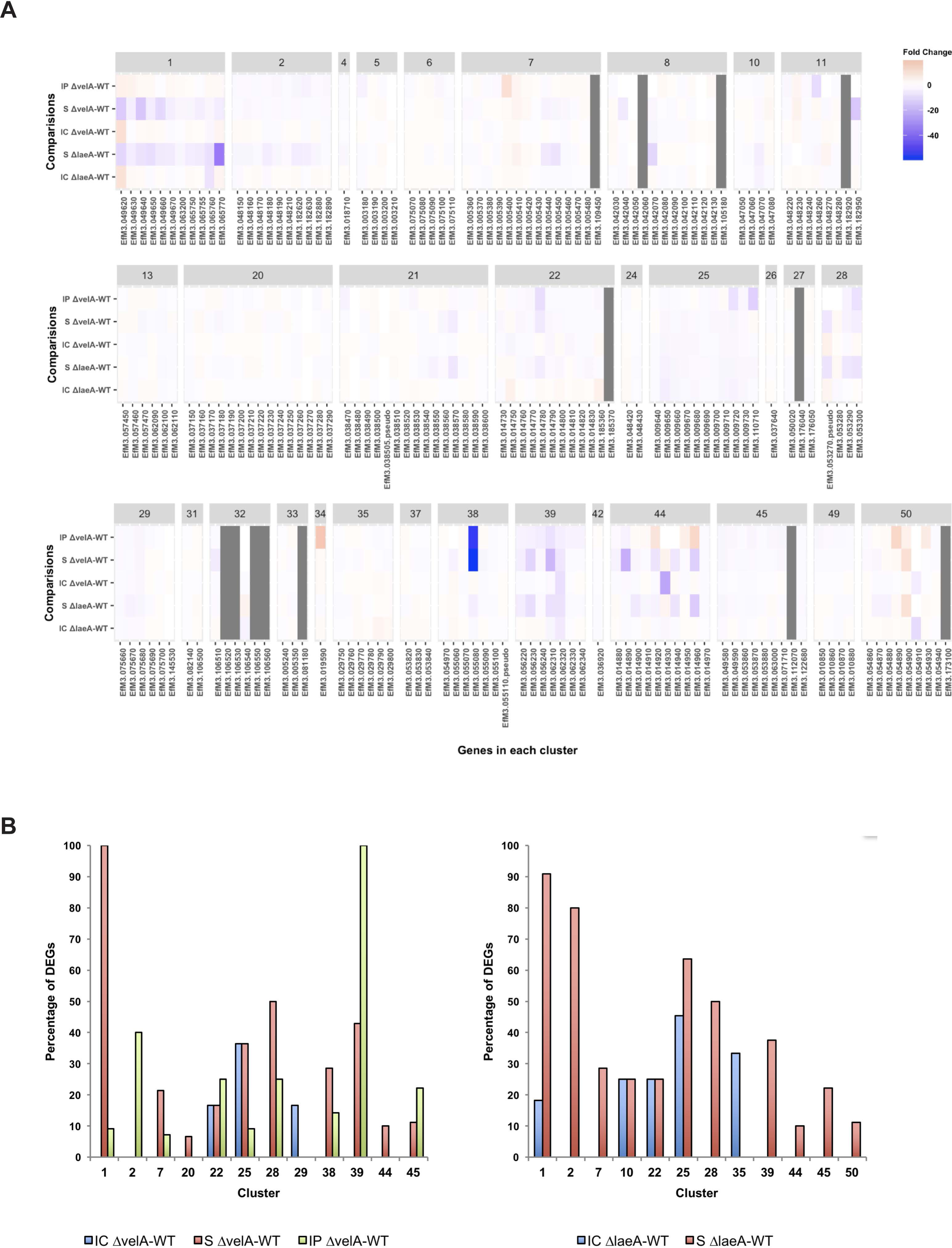
Transcriptomics profile of all secondary metabolism clusters in *Epichloë festucae*. a) Heatmap of all genes in each secondary metabolism cluster. Grey colour cells showing that in that condition none of the mutant or wild type are expressed. b) Percentage of DEGs in the clusters with at least one DEGs.

Our results show that VelA, similar to LaeA, is a key regulator for secondary metabolites gene expression and production in *E. festucae.* However, it seems VelA and LaeA exert different regulatory effects.

#### 5. Changes in expression of genes encoding putative small-secreted proteins

Microbes during infection of host plants produce effector proteins that are of key interest during plant-microbe interactions. Via a range of different molecular mechanisms these secreted proteins suppress or interfere with host immune responses or alter the host cell physiology, generating a favourable environment for infection and growth [59, 60]. Recently, 141 genes in *E. festucae* were reported as putative effectors [61]. Of these, 49 were differentially expressed in at least one of the comparisons described here (Fig. 6 and Table S7). Although the S Δ*vel*A-WT comparison had the highest number of SSPs (Fig. 6a) the IP Δ*vel*A-WT comparison had a much broader range of fold changes (72.4 and −99.8) (Fig. 6b). *velA* and *laeA* showed different regulatory effects on SSPs across the different conditions (Fig. 6c) with some being unique to particular conditions (Fig. 6d).

**Fig. 6.**
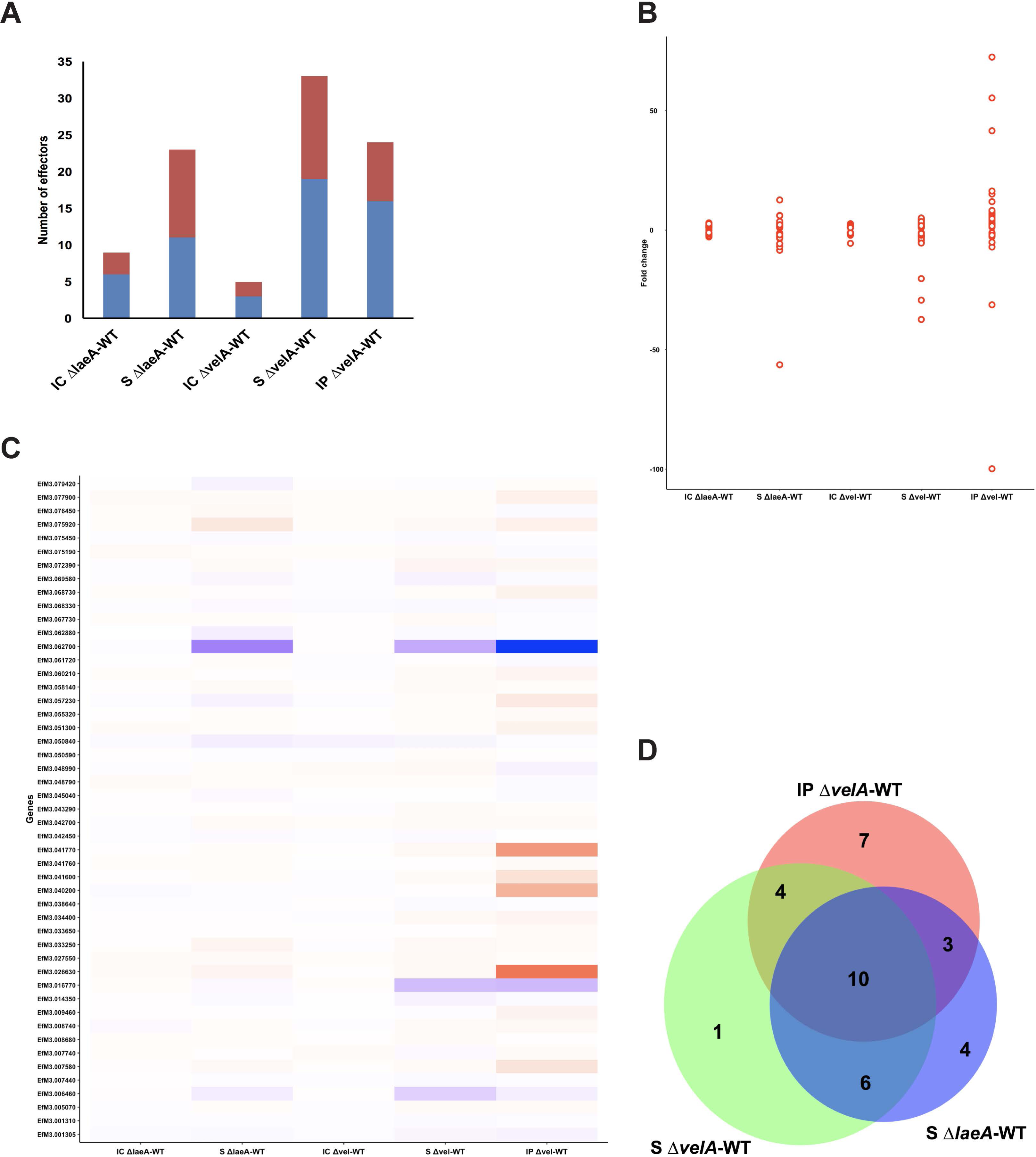
DEGs with putative SSP function. a) Number of DEGs, down or up regulated, in each comparison with putative SSPs functions. b) Distribution of SSPs fold change in different conditions, each point representing one SSP. c) Heat map showing the different ranges of regulatory effects of VelA and LaeA on the expression of putative SSPs in different conditions. d) Venn diagram shows the common SSPs between different comparisons which significantly differentially expressed.

Most of the predicted gene products have no homologues with annotated molecular function except four genes EfM3.001305, EfM3.001310, EfM3.055320 and EfM3.079420 (Table S7). EfM3.001305 and EfM3.001310 are homologues with a killer toxin gene, *kp4*, from *Metarhizium spp* that is a lethal protein via inhibition of calcium channels [62]. EfM3.055320 was only expressed in the in planta comparisons and is a homologue of Cu/Zn superoxide dismutase from *Claviceps purpurea* [6] which are involved in reducing superoxide radicals generated by host plant defence mechanisms. EfM3.079420 is a homologue of a fungal hydrophobin domain-containing protein in *Pochonia chlamydosporia* [63].

Three putative effectors, which are some of the core symbiosis genes [20] showed the highest differential gene expression and were down-regulated. EfM3.016770, EfM3.062700 and EfM3.014350 called *sspM, sspN* and *sspO*, respectively, were functionally studied in Hassing et al (2019) but no clear function was identified presumably due to redundancy functions of effectors. These 3 effector candidates were also down regulated in all S Δ*velA*-WT, IP Δ*velA*-WT and S Δ*laeA*-WT comparisons (Table S7).

Although only 27% (49 genes) of the Eaton et al symbiosis core genes are common with DEGs in the mature infected plants with Δ*velA*, 16% (8 genes) of these are SSPs, which are 57% (8/14) of the total SSP genes in the Eaton et al core set.

## Discussion

The *velA* gene is a well-known regulatory gene required for different fungal cellular and developmental functions, including secondary metabolism, pathogenicity, sexual and asexual development and fungal morphology and growth [25, 64]. In addition, we recently showed that it is required for the symbiotic interaction of *E. festucae* with perennial ryegrass [11]. Here, by using comparative transcriptomics, VelA regulatory effects on the *E. festucae* transcriptome profile growing in different conditions (in culture, inoculated seedlings and mature infected plants) were tested and the results compared with similar experiments, previously reported, on the other fungal global regulator LaeA [12, 37].

The numbers of DEGs in S Δ*vel*A-WT and IP Δ*velA*-WT comparisons were 5.1 and 3.9 times, higher than IC Δ*velA*-WT, respectively. This suggests that VelA has a stronger regulatory effect when growing in seedlings and mature plants under likely nutrient limited conditions, compared to PDA culture, a nutrient rich condition. This also correlates with our previous report that showed much higher expression of *velA* under nutrient limited medium (water agar) compared to nutrient rich medium (PDA) [11], which is similar to LaeA [12].

The small proportion of DEGs common between IC Δ*velA*-WT, S Δ*velA*-WT and IP Δ*velA*-WT comparisons suggests condition dependent regulatory roles of VelA on *E. festucae* gene expression, similar to LaeA. It is possible that VelA forms different protein complexes in different conditions as was shown in *A. nidulans* [25]. Other possibilities are different post-translational modifications or localisation under different conditions.

Compared to the core symbiotic fungal gene set identified by Eaton *et al.* (2015) only a small proportion of DEGs from both IP Δ*velA*-WT and S Δ*velA*-WT comparisons were in common. This may partly be a result of different tissue samples used for the transcriptomics analyses between the two studies. However, based on the difference in hyphal in planta characteristics between mutant *velA* and *proA, noxA* and *sakA* mutants, it is most likely VelA, like LaeA, regulates separate mechanisms important in mutualistic interactions than ProA, NoxA and SakA. One of the distinct differences between in planta fungal growth of *velA* and *laeA* mutants, compared to *proA, noxA* and *sakA* mutants was chitin distribution. In *velA* and *laeA* mutants, chitin distribution was very similar to the wild type interaction [11, 12], whereas for *proA, noxA* and *sakA* mutants, chitin distribution was increased [20].

Comparative transcriptomics studies of Δ*velA* mutants in different fungi show conserved regulatory roles of VelA on secondary metabolism, CAZyme biosynthesis, morphogenesis, development and cellular metabolism [27-33, 35], that are quite similar with some reported roles of LaeA [35, 65-67]. In the fungal transcriptomics analysis of S Δ*velA*-WT and IP Δ*velA*-WT and S Δ*laeA*-WT comparisons, up regulation was observed in the expression of genes that encode putative nutrient transporters, host cell wall degrading enzymes and small secreted proteins. In these three functional categories, DEGs in the IP Δ*velA*-WT comparison showed higher differential expression with more genes involved, compared with seedling comparisons, although total DEGs in this comparison was not higher than others. This suggests that VelA has stronger regulatory influence on fungal gene expression for these three functional categories during interaction with mature plants compared to the early stages of infection.

Fungal transporters of nitrate, ammonium, peptides and carboxylic acid were up-regulated in mature plants infected with the Δ*velA* mutant, while in seedlings inoculated with Δ*velA* and Δ*laeA* mutants peptide and ammonium transporters were up-regulated. This may suggest that in these associations, fungi are under starvation conditions, especially in mature infected plants as was previously suggested for *sakA, noxA* and *proA* mutants interactions [20]. The observed abnormal in planta hyphal growth of Δ*velA* mutant fungi and invasion of vascular bundles are a sign of starvation and these hyphae may be acting as a sink to absorb nutrients from host phloem [11]. Our *in vitro* analyses also show that Δ*velA* and Δ*laeA* mutants under starvation conditions grow abnormally but under rich conditions mostly grow normally [11, 12]. Also *in vitro* analyses of *velA* expression showed highest levels of their transcripts under starvation conditions [11]. Based on these observations, we speculated that in *E. festucae* during in planta growth, high expression of *velA* suppresses transporters and other starvation-response genes so that the fungus stops growing, leading to the growth restriction observed in wild type associations. Deleting *velA* in *E. festucae* changes its mutualistic interaction with ryegrass to a more antagonistic one and mutant fungi caused death in most inoculated plants. It has been shown that nitrogen metabolism generally plays a key role in pathogenicity [68, 69]. However, other than nitrogen transporters no predicted genes involved in nitrogen metabolism were found to be differentially expressed in this study. Possibly, changes in nitrogen transport may contribute to the host death, with increased fungal growth and insufficient nutrition for the seedling. This is supported by a comparative transcriptomics study of the *Botrytis cinerea* interaction with *Phaseolus vulgaris* in which it was suggested that up regulation of 2% of DEGs that encode sugar, amino acid and ammonium transporters and glycoside hydrolases is the reason for the virulence disability of the Δ*velA* mutant fungus [34]. It was also suggested that the transcriptomics pattern in this study is a sign of the Δ*velA* mutant response to nutrient starvation conditions [34].

In different transcriptomics analyses, VelA regulatory effects on secondary metabolite gene clusters were shown in different fungi [27-31, 33, 35]. In this study, we showed VelA positively regulates 48.4% of the gene clusters for secondary metabolism in *E. festucae* including the two well-known clusters for ergot alkaloids and indole diterpenes. The influence of VelA on ergot alkaloid gene expression in mature plants was much lower than in seedlings inoculated with Δ*velA* and in fact in mature plants only one gene is differentially expressed, being up-regulated compared with seedlings where most genes are down-regulated. VelA regulates indole diterpene gene expression only in infected mature plants, exerting a much weaker influence than seedlings inoculated with the Δ*laeA* mutant. Similar to VelA, NoxA and ProA were shown to have weak regulatory effects on ergot alkaloid and indole diterpene gene expression in infected mature plants, although SakA has a stronger positive regulatory effect [20]. The chemical concentration of ergot alkaloids in mature plants infected with the Δ*velA* mutant showed no difference to the wild type, correlating with the weak regulatory effect of VelA on their gene expression. The chemical concentration of most indole diterpenes was significantly greater in Δ*velA* mutant infected plants but this did not correlate with greater gene expression. It is likely that the higher biomass of Δ*velA* mutant *E. festucae* in infected plants, even led to the higher alkaloid levels. However, there is not always a strong correlation of gene expression with alkaloid production, as also observed in Δ*sakA*, Δ*noxA* and Δ*proA* mutant associations [20].

Most ergot alkaloids were detected at lower concentrations in Δ*laeA* mutant infected plants and this was less significant for Δ*velA* mutant infected plants compared to wild type. Despite this, this difference was greater for most of the indole-diterpene alkaloids. Therefore, VelA and LaeA exert different influences on alkaloid production in the *E. festucae*-perennial ryegrass association.

Of all the putative SSPs in *E. festucae*, 3.4%, 30% and 16.7% were differentially expressed in IC Δ*velA*-WT, S Δ*velA*-WT and IP Δ*velA*-WT comparisons, respectively. It seems VelA is a key regulator of SSPs expression in *E. festucae* during its interaction with ryegrass. The greater numbers of differentially expressed SSPs in infected seedlings and mature plants inoculated with the Δ*velA* mutant, compared to in culture conditions, demonstrates their importance during the infection process and survival. This has also been shown in other pathogenic or symbiotic fungi (Reviewed in Presti et al 2015). Although there are more differentially expressed SSPs in seedlings inoculated with the mutant compared to mature infected plants a much broader range of fold changes were observed in mature plants. This suggests condition-dependent regulatory roles of VelA for SSPs’ gene expression that possibly is a result of different mechanisms involved during the fungal interaction under these two conditions. Possibly, VelA by regulating different SSPs during the infection process and in mature infected plants positively regulates SSPs that are required for establishing and maintaining symbiotic interaction, or negatively regulates SSPs that lead to incompatible interactions. On the other hand, 6.2% and 16% of SSPs were differentially expressed in IC Δ*laeA*-WT and S Δ*laeA*-WT, respectively. Compared to the *velA* related comparisons, it seems LaeA has higher regulatory effects in culture and lower in seedlings compared to VelA. This suggests different regulatory effects for these two related global regulators.

In conclusion, this study provides a significant understanding on the strong regulatory effects of VelA on the *E. festucae* transcriptome profile during compatible and incompatible interactions with ryegrass. These regulatory effects could explain some of the observed phenotypes resulting from deleting *velA* in this fungus [11]. It seems that during the interaction of *E. festucae* with ryegrass, VelA controls fungal gene expression to supress or activate genes involved in either inducing or reducing the hostplant response.

## Methods

### Sample preparation

Total RNA was extracted from samples comprising fungus grown in culture, perennial ryegrass seedlings and mature perennial ryegrass plants (Table 1). For culture conditions, *E.* festucae wild type and Δ*velA* mutant strains were grown for two weeks on cellophane covered PDA medium in full darkness prior to harvest. Seedlings were grown and harvested as previously described [37]. 7-10 d old endophyte-free seedlings of perennial ryegrass (*L. perenne* ‘Nui’) were inoculated with wild type and Δ*velA* mutant strains *E.* festucae (Generated previously [11, 70]), which were grown for two weeks on PDA medium. Inoculated seedlings, grown for two weeks under 16 h of 650 W/m^2^ light and 8 h dark and after freezing, samples from 4 cm upwards and 0.5 cm downwards from the meristem were collected for RNA extraction and around 100 seedlings for each sample were pooled in three replicates for each treatment.

For the mature plants, the top 4 cm of the newest mature blade of infected plants with different strains were collected in three replicates (three mature plants infected with the same strain) for each treatment.

RNA quality and quantity were determined using an Agilent 2100 Bioanalyzer (Agilent Technologies), Nanodrop Lite spectrophotometer (Thermo scientific) and gel electrophoresis using a 1% agarose gel. RNA samples on dry ice were sent to the Beijing Genomics Institute (BGI, Hong Kong) for sequencing and 2 µg of RNA sample used to prepare libraries by BGI standard method (http://www.bgi.com/services/genomics/rna-seqtranscriptome/#tab-id-2). Samples were sequenced in two lanes of an Illumina HiSeq4000 (paired end, 100-bp reads).

### RNA extraction and quantitative real-time RT-PCR analysis

RNA extraction and complementary DNA (cDNA) were synthesised from 2 μg of RNA and Real time qPCR was performed with 1 μl of cDNA as described previously [12] using primers that amplified target genes (Table S1). RNA samples alone (no reverse transcription) and water-only (no template) controls were used to detect gDNA and experimental contamination. Comparative ΔCt normalized to gamma actin and 60S ribosomal protein L35 was used to calculate transcription levels by using the geNorm algorithm automated in CFX manager software (Bio-Rad).

### HiSeq results analysis

HiSeq results analysis were done as described in Rahnama *et al.* (2019) [37] by using a combination RNA-star version 2.5.0c [71], for mapping the HiSeq reads against the genome dataset of *E. festucae* Fl1, with EdgeR package version 3.10.5 [72], for counting the mapped genes. Fold changes and p-values were generated using Exact Tests for differences between two groups of Negative-Binomial Counts.

Gene ontology (GO) and functional annotation analysis of differentially expressed genes were performed as described in Rahnama *et al.* (2019) [37].

### General bioinformatics analyses

Venn diagrams were drawn using BioVenn online software [73]. Volcano plots were drawn in R statistical software environment version 3.2.1 [74].

## Acknowledgment

We thank C.R. Voisey, W.R. Simpson, W. Mace and A. deBonth for technical assistance (Forage Improvement, AgResearch Grasslands), and Biotelliga Ltd for providing laboratory space.

## Authors’ contributions

DJF, RDJ and MR designed the project. MR performed the experiments and drafted the manuscript. MR and PM performed bioinformatics. MR and RDJ revised the manuscript.

## Availability of data and materials

HiSeq Illumina sequencing included 36 raw sequence data sets that have been deposited into the NCBI SRA database with the BioProject ID PRJNA578737.

