## Supplemental Tables for "VelA and LaeA are key regulators of the *Epichloë festucae* transcriptomic response during symbiosis with perennial ryegrass"

**Table S1.** Primers for qRT-PCR used in this study.

| Name | Sequence (5'-3') | Purpose | Gene Model |
|---|---|---|---|
| LaeAq-F | GGTACCGGAATATGGGCCATT | *laeA* Fwd | EfM3.069170 |
| LaeAq-R | GAATGAGGGCTGGCTGAATC | *laeA* Rev | |
| VelAq2-F | GCTCAGGGATGAAGGCGAAT | *velA* Fwd | EfM3.049680 |
| VelAq3-R | TGGCGTTGTAGTCGAAAGTGA | *velA* Rev | |
| Actinq.IDT-F | GTCTTGAGTCTGGCGGTATC | *gamma actin* Fwd | EfM3.017740 |
| Actinq.IDT-R | CTTCTGCATACGGTCGGAG | *gamma actin* Rev | |
| 60SqIDT-F | GCAGCAAGTTGAACAAGATCC | 60S ribosomal protein L35 | EfM3.073270 |
| 60SqIDT-R | CACGGGTTTGCTTGACAC | 60S ribosomal protein L35 | |
| perA-F | TGACGGTTGAAGTATGGCTG | *perA* Fwd | EfM3.018710 |
| perA-R | TCTGTGAAGGTGTGGCATG | *perA* Rev | |
| ltmG-F | TGACCGACGCCATTAATGAG | *ltmG* Fwd | EfM3.182880 |
| ltmG-R | TCCACGTCGCAGTTTAGATG | *ltmG* Rev | |
| dmaWq-F | CCACTCGTACATTTCCTTCTC | *dmaW* Fwd | EfM3.065770 |
| dmaWq-R | TTCGGGAAACACAGGCCATTC | *dmaW* Rev | |

**Table S2-** General description of mRNA-sequencing results

| mRNA-sequencing reads | In culture | In seedling | In planta |
|---|---|---|---|
| Total read number | 1,132,721,194 | 905,625,652 | 810,694,678 |
| Number of total reads after quality trimming | 1,131,271,924 | 904,675,122 | 809,938,396 |
| Percentage of total reads after quality trimming | 99.87% | 99.90% | 99.91% |
| Number of mapped reads | 1,083,417,500 | 764,871,314 | 724,293,326 |
| Proportion of mapped reads (% of trimmed reads) | 95.77% | 84.55% | 89.43% |
| Number of reads mapped to Endophyte genome | 1,080,853,724 | 48,000,483 | 13,360,773 |
| Percentage of mapped fungal reads to total mapped reads | 99.76% | 6.28% | 1.84% |

**Table S3-** Percentage of DEGs with their primary functions onto 'Biological Process' Gene Ontology.

| Model | Biological process categories | Percentage of DEGs | | |
|---|---|---|---|---|
| | | **IC Δ*velA*-WT** | **S Δ*velA*-WT** | **IP Δ*velA*-WT** |
| GO:0032502 | **developmental process** | 0.00 | 0.00 | 0.44 |
| GO:0023052 | **signaling** | 1.75 | 1.71 | 1.33 |
| GO:0050789 | **regulation of biological process** | 8.77 | 4.44 | 5.33 |
| GO:0000003 | **reproduction** | 1.75 | 0.34 | 0.44 |
| GO:0016043 | **cellular component organization** | 0.00 | 0.34 | 3.11 |
| GO:0071554 | **cell wall organization or biogenesis** | 1.75 | 1.02 | 1.33 |
| GO:0065007 | **biological regulation** | 10.53 | 5.12 | 6.67 |
| GO:0048518 | **positive regulation of biological process** | 1.75 | 1.02 | 1.78 |
| GO:0048519 | **negative regulation of biological process** | 3.51 | 0.68 | 0.89 |
| GO:0019740 | **nitrogen utilization** | 1.75 | 0.68 | 0.44 |
| GO:0051704 | **multi-organism process** | 1.75 | 1.71 | 1.78 |
| GO:0009987 | **cellular process** | 28.07 | 18.43 | 25.33 |
| GO:0022414 | **reproductive process** | 1.75 | 0.34 | 0.44 |
| GO:0023046 | **signaling process** | 1.75 | 1.37 | 0.89 |
| GO:0008152 | **metabolic process** | 33.33 | 38.91 | 43.11 |
| GO:0051234 | **establishment of localization** | 8.77 | 6.48 | 9.78 |
| GO:0051179 | **localization** | 10.53 | 6.83 | 10.22 |
| GO:0040007 | **growth** | 0.00 | 0.34 | 0.44 |
| GO:0050896 | **response to stimulus** | 1.75 | 3.07 | 3.11 |
| GO:0044085 | **cellular component biogenesis** | 1.75 | 0.68 | 1.33 |
| | **No GO** | 68.42 | 57.00 | 61.33 |

**Table S4-** Differential expression of different velvet component in different comparisons. Fold changes show in bold are statistically significant (FDR≤0.05) changed more than two times.

| | **IC Δ*laeA*-WT** | **S Δ*laeA*-WT** | **IC Δ*velA*-WT** | **S Δ*velA*-WT** | **IP Δ*velA*-WT** |
|---|---|---|---|---|---|
| ***velD*** | 1.12 | 2.00 | -1.03 | 2.24 | -1.36 |
| ***velC*** | 1.40 | 1.58 | 1.42 | -1.19 | 2.55 |
| ***velB*** | -1.14 | 1.17 | -1.11 | 1.33 | -1.05 |
| ***velA*** | 1.01 | 1.03 | **-3.20** | **-3.46** | **-4.69** |
| ***laeA*** | **-3.50** | **-2.13** | **2.22** | **2.48** | 1.07 |

**Table S5-** Homologues of known fungal plant cell wall degrading enzymes in *E. festucae*

| Model | Family | E-value | Category | Gene Family |
|---|---|---|---|---|
| EfM3.016930 | 1 | 3.00E-23 | Accessary - cellulose | Alpha-glucosidase (Type 1) |
| EfM3.024120 | 1 | 0 | Accessary - cellulose | Alpha-glucosidase (Type 1) |
| EfM3.050100 | 1 | 1.00E-25 | Accessary - cellulose | Alpha-glucosidase (Type 1) |
| EfM3.022450 | 2 | 0 | Accessary - cellulose | Alpha-glucosidase (Type 2) |
| EfM3.049570 | 2 | 5.00E-17 | Accessary - cellulose | Alpha-glucosidase (Type 2) |
| EfM3.010800 | 3 | 0 | Accessary - pectin | Beta-D-galactosidase (Type 1) |
| EfM3.010800 | 3 | 0 | Accessary - pectin | Beta-D-galactosidase (Type 1) |
| EfM3.053570 | 3 | 3.00E-20 | Accessary - pectin | Beta-D-galactosidase (Type 1) |
| EfM3.075210 | 4 | 0 | Accessary - pectin | Beta-D-galactosidase (Type 2) |
| EfM3.008730 | 6 | 5.00E-106 | Accessary - pectin | Pectin methylesterase |
| EfM3.008610 | 9 | 7.00E-81 | Leaf Surface - Cutin | Cutinase |
| EfM3.066110 | 9 | 4.00E-60 | Leaf Surface - Cutin | Cutinase |
| EfM3.030520 | 10 | 2.00E-12 | Main-chain degrading - cellulose | Cellobiohydrolase (Type 1) |
| EfM3.032730 | 12 | 0 | Main-chain degrading - glacto(gluco)mannan | Alpha-mannosidase (Type 1) |
| EfM3.039340 | 12 | 3.00E-63 | Main-chain degrading - glacto(gluco)mannan | Alpha-mannosidase (Type 1) |
| EfM3.042900 | 12 | 0 | Main-chain degrading - glacto(gluco)mannan | Alpha-mannosidase (Type 1) |
| EfM3.044230 | 12 | 0 | Main-chain degrading - glacto(gluco)mannan | Alpha-mannosidase (Type 1) |
| EfM3.073730 | 12 | 0 | Main-chain degrading - glacto(gluco)mannan | Alpha-mannosidase (Type 1) |
| EfM3.078610 | 12 | 0 | Main-chain degrading - glacto(gluco)mannan | Alpha-mannosidase (Type 1) |
| EfM3.079010 | 12 | 2.00E-29 | Main-chain degrading - glacto(gluco)mannan | Alpha-mannosidase (Type 1) |
| EfM3.015680 | 13 | 0 | Main-chain degrading - glacto(gluco)mannan | Alpha-mannosidase (Type 2) |
| EfM3.020350 | 15 | 0 | Main-chain degrading - glacto(gluco)mannan | Beta-mannosidase |
| EfM3.056680 | 19 | 2.00E-27 | Main-chain degrading - pectin | Polygalacturonase |
| EfM3.061190 | 19 | 6.00E-85 | Main-chain degrading - pectin | Polygalacturonase |
| EfM3.040190 | 21 | 6.00E-98 | Main-chain degrading - xylan | Endoxylanase (Type 1) |
| EfM3.050060 | 22 | 1.00E-62 | Main-chain degrading - xylan | Endoxylanase (Type 2) |

**Table S6-** Differentially expressed genes of secondary metabolism gene clusters in different comparisons. Fold changes show in bold are statistically significant (FDR≤0.05) changed more than two times. Empty fold change cells are representative of not expressed gene in both Δ*laeA* and wild type strains. Clusters highlighted in grey colour are including DEGs. Alk: alkaloids, K: polyketide synthase, N: nonribosomal peptide synthetase, D: DMATS-family prenyltransferase, T: terpene cyclase, Misc.: miscellaneous.

| | | | Fold change | | | | |
|---|---|---|---|---|---|---|---|
| | Cluster | Cluster Type[a] | IC Δ*laeA*-WT | S Δ*laeA*-WT | IC Δ*velA*-WT | S Δ*velA*-WT | IP Δ*velA*-WT |
| EfM3.049620 | 1 | Alk, D, N | **9.2** | **-10.3** | **9.7** | **-13.4** | **4.1** |
| EfM3.049630 | 1 | Alk, D, N | **2.1** | **-4.8** | -1.1 | **-4.2** | 4.0 |
| EfM3.049640 | 1 | Alk, D, N | 1.8 | -7.3 | 1.9 | **-14.5** | 2.4 |
| EfM3.049650 | 1 | Alk, D, N | -1.2 | **-8.4** | 2.3 | **-4.6** | 2.7 |
| EfM3.049660 | 1 | Alk, D, N | 1.2 | **-5.6** | 1.4 | **-9.7** | -1.0 |
| EfM3.049670 | 1 | Alk, D, N | -1.2 | **-5.0** | -1.3 | **-4.1** | 2.6 |
| EfM3.063200 | 1 | Alk, D, N | -1.0 | **-2.7** | -1.1 | **-2.3** | 1.5 |
| EfM3.065750 | 1 | Alk, D, N | 1.5 | **-4.8** | 1.3 | **-2.5** | 1.1 |
| EfM3.065755 | 1 | Alk, D, N | 1.2 | **-5.3** | 1.0 | **-4.1** | -1.0 |
| EfM3.065760 | 1 | Alk, D, N | -7.0 | **-7.8** | -1.9 | **-2.3** | 2.3 |
| EfM3.065770 | 1 | Alk, D, N | 3.4 | **-34.7** | 3.5 | **-5.0** | 2.0 |
| EfM3.048150 | 2 | Alk, T, D | -1.7 | **-2.5** | 1.7 | -1.5 | -1.8 |
| EfM3.048160 | 2 | Alk, T, D | 1.2 | **-2.7** | 1.3 | -1.8 | -1.7 |
| EfM3.048170 | 2 | Alk, T, D | 1.3 | **-2.0** | 1.1 | -1.8 | **-2.0** |
| EfM3.048180 | 2 | Alk, T, D | -1.6 | **-4.4** | -1.5 | -1.5 | -1.6 |
| EfM3.048190 | 2 | Alk, T, D | 1.0 | **-2.5** | -1.2 | -1.7 | **-2.1** |
| EfM3.048210 | 2 | Alk, T, D | 1.1 | -1.6 | 1.1 | -1.4 | -1.5 |
| EfM3.182620 | 2 | Alk, T, D | 1.7 | **-4.8** | 1.2 | -1.8 | **-2.7** |
| EfM3.182630 | 2 | Alk, T, D | -1.0 | **-3.9** | -1.3 | -1.5 | **-3.1** |
| EfM3.182880 | 2 | Alk, T, D | 1.4 | **-2.8** | 1.2 | -1.3 | -1.7 |
| EfM3.182890 | 2 | Alk, T, D | 1.2 | -1.2 | -1.4 | -1.6 | -1.9 |
| EfM3.018710 | 4 | Alk, N | 1.0 | **-1.9** | 1.1 | -1.3 | -1.4 |
| EfM3.003180 | 5 | K | -1.1 | -1.2 | 1.0 | -1.0 | -3.2 |
| EfM3.003190 | 5 | K | 1.0 | -1.0 | -1.1 | 1.2 | 1.1 |
| EfM3.003200 | 5 | K | 1.2 | 1.8 | -1.0 | 1.6 | -1.2 |
| EfM3.003210 | 5 | K | 0.0 | 0.0 | 1.0 | 0.0 | 0.0 |
| EfM3.075070 | 6 | K | 1.0 | 1.1 | 1.0 | -1.0 | -1.4 |
| EfM3.075080 | 6 | K | 1.1 | 1.1 | 1.1 | 1.0 | 1.1 |
| EfM3.075090 | 6 | K | 1.1 | -1.2 | 1.0 | 1.2 | 1.4 |

| | | | | | | | |
|---|---|---|---|---|---|---|---|
| EfM3.075100 | 6 | K | 1.2 | 1.1 | 1.0 | 1.3 | -2.3 |
| EfM3.075110 | 6 | K | 1.1 | -1.2 | -1.0 | -1.5 | -1.0 |
| EfM3.005360 | 7 | K | 1.1 | -1.5 | -1.1 | -1.7 | -1.1 |
| EfM3.005370 | 7 | K | -1.1 | -1.2 | 1.0 | -1.5 | 1.6 |
| EfM3.005380 | 7 | K | -1.5 | 1.0 | -1.1 | 1.1 | -1.2 |
| EfM3.005390 | 7 | K | 1.0 | -1.2 | -1.0 | 1.1 | -1.7 |
| EfM3.005400 | 7 | K | 1.4 | 2.2 | 1.3 | 1.7 | **10.1** |
| EfM3.005410 | 7 | K | 1.7 | **3.8** | 1.2 | 3.5 | 3.2 |
| EfM3.005420 | 7 | K | 1.1 | 1.9 | -1.0 | **2.3** | 1.5 |
| EfM3.005430 | 7 | K | -1.1 | 1.2 | -1.1 | -1.0 | 1.1 |
| EfM3.005440 | 7 | K | -1.5 | **-5.4** | 1.0 | **-2.8** | -1.3 |
| EfM3.005450 | 7 | K | -1.4 | **-6.9** | -1.1 | **-2.5** | 1.1 |
| EfM3.005460 | 7 | K | -1.3 | **-2.1** | -1.2 | -1.9 | 1.7 |
| EfM3.005470 | 7 | K | -1.0 | -1.2 | 1.0 | -1.3 | 1.3 |
| EfM3.005480 | 7 | K | -1.1 | 1.0 | -1.1 | 1.2 | 1.4 |
| EfM3.109450 | 7 | K | | | | | |
| EfM3.042030 | 8 | K | -1.0 | 1.8 | 2.0 | 1.5 | -1.3 |
| EfM3.042040 | 8 | K | -1.5 | -1.1 | -1.1 | -5.8 | 0.0 |
| EfM3.042050 | 8 | K | 1.3 | 0.0 | 1.2 | 0.0 | 0.0 |
| EfM3.042060 | 8 | K | | | | | |
| EfM3.042070 | 8 | K | 1.4 | -10.0 | -2.7 | -2.3 | 0.0 |
| EfM3.042080 | 8 | K | -1.1 | 1.1 | -1.4 | -1.1 | 2.1 |
| EfM3.042090 | 8 | K | -1.0 | 1.0 | -1.3 | -1.5 | 1.0 |
| EfM3.042100 | 8 | K | -1.1 | 1.0 | -1.2 | 1.2 | -1.2 |
| EfM3.042110 | 8 | K | 1.8 | -1.1 | 1.3 | -1.6 | 1.2 |
| EfM3.042120 | 8 | K | 0.0 | 0.0 | 1.0 | 0.0 | 0.0 |
| EfM3.042130 | 8 | K | 1.5 | 1.1 | 1.3 | -1.1 | 2.0 |
| EfM3.105180 | 8 | K | | | | | |
| EfM3.047050 | 10 | K | -1.0 | -1.1 | -1.0 | 1.1 | -1.0 |
| EfM3.047060 | 10 | K | -1.0 | -1.1 | 1.0 | 1.0 | -1.0 |
| EfM3.047070 | 10 | K | **-3.0** | **-4.1** | -1.3 | -1.7 | -1.2 |
| EfM3.047080 | 10 | K | -1.6 | 1.6 | 1.2 | 2.8 | 1.7 |
| EfM3.048220 | 11 | K | 1.1 | -1.1 | 1.0 | -1.2 | -1.9 |
| EfM3.048230 | 11 | K | 1.5 | 0.0 | 2.5 | 4.0 | 3.0 |
| EfM3.048240 | 11 | K | 1.1 | -1.2 | -1.1 | -1.3 | -1.6 |
| EfM3.048260 | 11 | K | 1.4 | -1.9 | -1.4 | 2.3 | **-8.9** |
| EfM3.048270 | 11 | K | -1.4 | 3.3 | 1.8 | -2.0 | 0.0 |
| EfM3.048280 | 11 | K | -1.2 | -1.1 | 1.1 | 1.4 | -1.3 |
| EfM3.182920 | 11 | K | | | | | |
| EfM3.182950 | 11 | K | 1.2 | -1.7 | -1.7 | **-14.0** | 0.0 |
| EfM3.057450 | 13 | K | 1.1 | 1.0 | -1.0 | 1.1 | -1.0 |
| EfM3.057460 | 13 | K | -1.1 | -1.3 | -1.4 | 1.3 | 1.2 |
| EfM3.057470 | 13 | K | 1.3 | -1.1 | -1.5 | -1.9 | 1.2 |

| | | | | | | | |
|---|---|---|---|---|---|---|---|
| EfM3.062090 | 13 | K | 1.3 | 1.0 | 1.0 | -1.4 | 1.3 |
| EfM3.062100 | 13 | K | -1.2 | -1.7 | -1.4 | -1.8 | -1.5 |
| EfM3.062110 | 13 | K | -1.1 | -1.2 | -1.2 | 1.2 | -1.0 |
| EfM3.037150 | 20 | K-N | -1.3 | -1.2 | -1.0 | -1.3 | -1.1 |
| EfM3.037160 | 20 | K-N | -1.0 | 1.7 | 1.1 | **2.1** | -1.0 |
| EfM3.037170 | 20 | K-N | 1.1 | -1.0 | 1.1 | -1.1 | 1.2 |
| EfM3.037180 | 20 | K-N | 1.0 | -1.2 | -1.0 | -1.9 | 1.6 |
| EfM3.037190 | 20 | K-N | -1.1 | -1.3 | 1.0 | -1.2 | -1.4 |
| EfM3.037200 | 20 | K-N | -1.1 | -1.1 | 1.1 | -1.0 | -1.1 |
| EfM3.037210 | 20 | K-N | -1.0 | 1.6 | 1.0 | 1.2 | -1.5 |
| EfM3.037220 | 20 | K-N | -1.0 | 1.3 | 1.1 | 1.5 | -1.1 |
| EfM3.037230 | 20 | K-N | -1.0 | -1.1 | -1.0 | -1.3 | 1.4 |
| EfM3.037240 | 20 | K-N | -1.1 | 1.0 | 1.0 | -1.2 | 1.2 |
| EfM3.037250 | 20 | K-N | -1.1 | -1.1 | -1.0 | 1.2 | -1.1 |
| EfM3.037260 | 20 | K-N | 1.0 | -1.1 | -1.2 | -1.1 | -1.1 |
| EfM3.037270 | 20 | K-N | 0.0 | 0.0 | 1.0 | 0.0 | 0.0 |
| EfM3.037280 | 20 | K-N | 1.6 | -1.3 | 1.0 | -1.1 | -2.0 |
| EfM3.037290 | 20 | K-N | -1.0 | -1.2 | -1.0 | -1.2 | 1.2 |
| EfM3.038470 | 21 | K, D | 1.1 | 1.3 | 1.1 | -1.5 | -3.0 |
| EfM3.038480 | 21 | K, D | 0.0 | 0.0 | 1.0 | 0.0 | 0.0 |
| EfM3.038490 | 21 | K, D | 1.0 | -1.2 | -1.0 | -1.3 | 1.2 |
| EfM3.038500 | 21 | K, D | -1.0 | 1.4 | -1.1 | 1.8 | 1.3 |
| EfM3.038505.<br>pseudo | 21 | K, D | 0.0 | 0.0 | 1.0 | 0.0 | 0.0 |
| EfM3.038510 | 21 | K, D | -1.0 | 1.4 | 1.0 | 1.5 | 1.1 |
| EfM3.038520 | 21 | K, D | 1.0 | 1.0 | 1.0 | 1.1 | -1.1 |
| EfM3.038530 | 21 | K, D | 1.4 | -1.0 | 1.1 | -1.1 | -1.9 |
| EfM3.038540 | 21 | K, D | 1.2 | -2.1 | -1.6 | -1.6 | 1.0 |
| EfM3.038550 | 21 | K, D | 1.1 | -3.0 | -1.2 | 1.1 | 1.0 |
| EfM3.038560 | 21 | K, D | -1.1 | -1.6 | -1.0 | -2.5 | 1.3 |
| EfM3.038570 | 21 | K, D | -1.1 | -6.0 | -1.2 | -4.8 | -1.7 |
| EfM3.038580 | 21 | K, D | 1.0 | -1.1 | 1.1 | -1.0 | -1.2 |
| EfM3.038590 | 21 | K, D | 1.1 | 1.1 | 1.0 | 1.0 | -1.2 |
| EfM3.038600 | 21 | K, D | -1.1 | -1.0 | 1.2 | -1.1 | 1.4 |
| EfM3.014730 | 22 | K, K | -1.2 | 1.3 | -1.2 | -2.0 | 1.4 |
| EfM3.014750 | 22 | K, K | **4.2** | 1.6 | -2.6 | 1.3 | -1.2 |
| EfM3.014760 | 22 | K, K | 1.0 | **-2.1** | -1.5 | **-2.1** | -1.7 |
| EfM3.014770 | 22 | K, K | -1.0 | -2.1 | -1.6 | -2.0 | **-2.1** |
| EfM3.014780 | 22 | K, K | 1.4 | **-4.4** | 1.2 | **-5.5** | **-8.5** |
| EfM3.014790 | 22 | K, K | 1.2 | -1.8 | **-2.1** | -1.9 | -1.4 |
| EfM3.014800 | 22 | K, K | **2.4** | -1.9 | 1.6 | -1.7 | -1.4 |
| EfM3.014810 | 22 | K, K | 1.1 | 1.1 | **-2.1** | -1.1 | -1.4 |
| EfM3.014820 | 22 | K, K | 1.7 | -1.4 | -1.4 | -1.3 | -1.5 |

| | | | | | | | |
|---|---|---|---|---|---|---|---|
| EfM3.014830 | 22 | K, K | 1.2 | **-2.1** | -1.1 | -1.4 | -1.2 |
| EfM3.185360 | 22 | K, K | **4.7** | -1.3 | 2.6 | -1.2 | **-2.1** |
| EfM3.185370 | 22 | K, K | | | | | |
| EfM3.048420 | 24 | N | 1.5 | 1.3 | 1.0 | -1.1 | -1.1 |
| EfM3.048430 | 24 | N | -1.0 | -1.4 | -1.0 | -1.1 | 1.0 |
| EfM3.009640 | 25 | N | -1.2 | -1.6 | -1.2 | 1.1 | -1.0 |
| EfM3.009650 | 25 | N | **-4.0** | **-3.5** | **-2.4** | **-2.2** | -2.0 |
| EfM3.009660 | 25 | N | **-2.6** | **-2.7** | **-2.2** | **-2.0** | -1.1 |
| EfM3.009670 | 25 | N | **-2.0** | **-2.2** | -1.9 | -1.8 | -1.1 |
| EfM3.009680 | 25 | N | -1.9 | -1.8 | -1.9 | -1.7 | 1.2 |
| EfM3.009690 | 25 | N | -1.4 | -1.6 | -1.2 | -1.6 | -1.4 |
| EfM3.009700 | 25 | N | -1.4 | **-2.1** | -1.5 | **-2.2** | -1.8 |
| EfM3.009710 | 25 | N | **-2.7** | **-2.5** | **-2.3** | -1.9 | 1.6 |
| EfM3.009720 | 25 | N | **-3.1** | **-2.6** | **-2.3** | **-2.4** | -7.7 |
| EfM3.009730 | 25 | N | -2.0 | **-2.2** | -1.7 | -1.8 | **-2.1** |
| EfM3.110710 | 25 | N | -1.1 | -2.2 | -1.2 | 1.3 | -11.8 |
| EfM3.037640 | 26 | N | 1.0 | -1.1 | 1.1 | -1.1 | -1.1 |
| EfM3.050020 | 27 | N | -1.2 | -1.1 | 1.0 | -1.1 | 1.2 |
| EfM3.176040 | 27 | N | | | | | |
| EfM3.176050 | 27 | N | -1.2 | -1.1 | 1.1 | 1.4 | 1.2 |
| EfM3.053270.<br>pseudo | 28 | N | 3.2 | -6.8 | 4.9 | -6.2 | 0.0 |
| EfM3.053280 | 28 | N | -1.8 | 2.5 | 1.0 | 2.1 | 0.0 |
| EfM3.053290 | 28 | N | -1.1 | **-5.9** | -1.3 | **-2.9** | -3.0 |
| EfM3.053300 | 28 | N | 1.1 | **-6.9** | 1.8 | **-4.5** | **-7.5** |
| EfM3.075660 | 29 | N | -1.4 | -2.1 | -1.2 | -1.6 | -1.7 |
| EfM3.075670 | 29 | N | -1.8 | -3.2 | -1.8 | -1.5 | -2.1 |
| EfM3.075680 | 29 | N | -1.8 | -2.6 | **-2.1** | -2.4 | -2.4 |
| EfM3.075690 | 29 | N | 1.1 | -1.2 | 1.0 | 1.0 | -1.5 |
| EfM3.075700 | 29 | N | 1.1 | 1.2 | -1.1 | 1.2 | -1.0 |
| EfM3.145530 | 29 | N | 0.0 | 0.0 | 1.0 | 0.0 | 0.0 |
| EfM3.082140 | 31 | N | -1.0 | -1.1 | -1.0 | 1.2 | -1.1 |
| EfM3.106500 | 31 | N | 1.1 | 1.3 | -1.1 | 1.1 | 1.1 |
| EfM3.106510 | 32 | N | 1.6 | -1.8 | -1.2 | -5.2 | -1.5 |
| EfM3.106520 | 32 | N | | | | | |
| EfM3.106530 | 32 | N | | | | | |
| EfM3.106540 | 32 | N | -3.3 | 3.7 | -1.3 | 0.0 | 0.0 |
| EfM3.106550 | 32 | N | | | | | |
| EfM3.106560 | 32 | N | | | | | |
| EfM3.005240 | 33 | N | -1.0 | 1.1 | -1.1 | 1.2 | -1.1 |
| EfM3.005350 | 33 | N | 1.0 | -1.5 | -1.0 | -1.2 | 1.9 |
| EfM3.081180 | 33 | N | | | | | |
| EfM3.019590 | 34 | N | 1.4 | -1.3 | 1.8 | 1.9 | 17.9 |

| | | | | | | | |
|---|---|---|---|---|---|---|---|
| EfM3.029750 | 35 | N | -1.0 | -1.4 | -1.0 | -1.3 | -1.0 |
| EfM3.029760 | 35 | N | -1.1 | -1.1 | 1.0 | -1.0 | 1.1 |
| EfM3.029770 | 35 | N | 1.4 | 1.3 | 1.2 | -1.4 | 1.1 |
| EfM3.029780 | 35 | N | 1.1 | -1.1 | 1.0 | 1.0 | -1.1 |
| EfM3.029790 | 35 | N | **3.3** | 1.1 | 1.4 | -1.1 | -1.3 |
| EfM3.029800 | 35 | N | **2.4** | 1.1 | 1.1 | -1.5 | -1.1 |
| EfM3.053820 | 37 | N | 1.2 | -1.2 | 1.0 | 1.2 | 1.3 |
| EfM3.053830 | 37 | N | 1.2 | 1.5 | -1.1 | -1.3 | -1.1 |
| EfM3.053840 | 37 | N | 1.2 | 1.1 | -1.0 | -1.1 | -1.1 |
| EfM3.054970 | 38 | N | -1.1 | 1.2 | -1.1 | -1.0 | 1.2 |
| EfM3.055060 | 38 | N | 2.5 | 1.6 | 2.5 | -3.5 | 0.0 |
| EfM3.055070 | 38 | N | 1.0 | -1.0 | -1.1 | 1.1 | 1.8 |
| EfM3.055080 | 38 | N | -2.0 | -3.9 | 1.3 | **-58.0** | **-55.3** |
| EfM3.055090 | 38 | N | 1.1 | -1.2 | -1.1 | **-2.7** | 1.3 |
| EfM3.055100 | 38 | N | -1.1 | -1.2 | -1.1 | -1.2 | -1.1 |
| EfM3.055110.pseudo | 38 | N | -1.0 | 1.1 | 1.0 | -1.0 | -1.0 |
| EfM3.056220 | 39 | N | 1.2 | **-4.8** | -1.1 | **-4.2** | **-3.4** |
| EfM3.056230 | 39 | N | 1.3 | **-8.4** | -1.3 | **-10.7** | **-3.8** |
| EfM3.056240 | 39 | N | 1.3 | -1.7 | -1.0 | -2.1 | **-4.7** |
| EfM3.062310 | 39 | N | -2.8 | **-7.5** | -1.7 | **-18.5** | **-7.5** |
| EfM3.062320 | 39 | N | -5.4 | -7.4 | -8.2 | -6.0 | **-4.2** |
| EfM3.062330 | 39 | N | -1.0 | -1.4 | -1.0 | -1.2 | **-3.0** |
| EfM3.062340 | 39 | N | 1.0 | -1.4 | 1.0 | -1.3 | **-3.2** |
| EfM3.036920 | 42 | N | -1.2 | -1.2 | -1.1 | -1.0 | 1.1 |
| EfM3.014880 | 44 | N, D | 1.2 | 1.7 | -1.1 | -3.5 | -1.7 |
| EfM3.014890 | 44 | N, D | -1.2 | -1.4 | -1.7 | -20.7 | -3.5 |
| EfM3.014900 | 44 | N, D | -1.1 | -5.5 | -1.4 | -2.4 | 1.3 |
| EfM3.014910 | 44 | N, D | 1.3 | -2.5 | 1.0 | -1.4 | 2.0 |
| EfM3.014920 | 44 | N, D | -1.6 | **-3.2** | -1.6 | **-6.9** | 8.7 |
| EfM3.014930 | 44 | N, D | -1.9 | 3.6 | -23.3 | 4.0 | 0.0 |
| EfM3.014940 | 44 | N, D | -1.2 | -6.8 | 1.6 | -6.2 | 0.0 |
| EfM3.014950 | 44 | N, D | 1.1 | 1.2 | -1.4 | -1.5 | 5.1 |
| EfM3.014960 | 44 | N, D | 1.0 | -13.2 | 1.3 | -12.0 | 12.9 |
| EfM3.014970 | 44 | N, D | -1.0 | -1.4 | -1.3 | -1.4 | -1.5 |
| EfM3.049580 | 45 | T | -1.6 | **-2.4** | -1.3 | **-2.8** | **-2.4** |
| EfM3.049590 | 45 | T | 1.1 | -1.8 | -1.1 | -1.8 | -2.1 |
| EfM3.053860 | 45 | T | -1.3 | -1.5 | -1.3 | -2.0 | -1.3 |
| EfM3.053870 | 45 | T | -1.1 | **-2.1** | -1.2 | -1.7 | **-2.3** |
| EfM3.053880 | 45 | T | 1.0 | -1.3 | -1.0 | -1.3 | -1.1 |
| EfM3.063000 | 45 | T | -1.2 | 1.3 | -1.1 | -1.5 | -1.9 |
| EfM3.071710 | 45 | T | 1.0 | -1.5 | 1.0 | -1.9 | -2.7 |
| EfM3.112070 | 45 | T | | | | | |

| | | | | | | | |
|---|---|---|---|---|---|---|---|
| EfM3.122680 | 45 | T | -1.1 | -1.6 | -1.1 | -1.5 | -1.2 |
| EfM3.010850 | 49 | Misc. | 1.2 | 1.1 | -1.1 | -1.3 | -1.9 |
| EfM3.010860 | 49 | Misc. | -1.0 | 1.1 | -1.0 | -1.3 | -1.6 |
| EfM3.010870 | 49 | Misc. | -1.1 | -1.3 | -1.0 | -1.2 | -2.0 |
| EfM3.010880 | 49 | Misc. | 1.1 | -1.1 | 1.0 | -1.2 | -1.5 |
| EfM3.054860 | 50 | Misc. | 1.1 | 1.1 | 1.1 | -1.7 | -1.1 |
| EfM3.054870 | 50 | Misc. | 1.0 | **-2.9** | -1.2 | -2.1 | 1.1 |
| EfM3.054880 | 50 | Misc. | 1.7 | -2.9 | -1.1 | 1.2 | 1.5 |
| EfM3.054890 | 50 | Misc. | 1.2 | 2.0 | -1.0 | 2.1 | 12.9 |
| EfM3.054900 | 50 | Misc. | 1.4 | 9.3 | 1.5 | 9.8 | 4.9 |
| EfM3.054910 | 50 | Misc. | -9.1 | 0.0 | -4.7 | 0.0 | 0.0 |
| EfM3.054930 | 50 | Misc. | 1.2 | 1.5 | -1.0 | -1.0 | 6.2 |
| EfM3.054940 | 50 | Misc. | -1.0 | 1.0 | 1.0 | -1.2 | -1.2 |
| EfM3.173100 | 50 | Misc. | | | | | |

**Table S7-** Putative small secreted proteins that differentially expressed in at least one of comparisons. Bold numbers are the fold changes that are significantly (FDR ≤ 0.05) changed more than two times.

| Gene | Name | IC Δ*laeA*-WT | S Δ*laeA*-WT | IC Δ*velA*-WT | S Δ*velA*-WT | IP Δ*velA*-WT | Presence in core set | Blast accession number | blast functions |
|---|---|---|---|---|---|---|---|---|---|
| EfM3.001305 | - | **-2.2** | -3.5 | -1.5 | **-4.1** | -5.16 | No | KID97701 | Killer toxin, Kp4, partial [Metarhizium majus ARSEF 297] |
| EfM3.001310 | - | -1.6 | -1.4 | -1.3 | **-2.7** | -1.54 | No | XP_014544272 | Calcium Channel Inhibitor, partial [Metarhizium brunneum ARSEF 3297] |
| EfM3.005070 | - | -1.2 | 1.3 | -1.2 | **2.4** | **3.11** | No | XP_018136637 | cupin superfamily protein [Pochonia chlamydosporia 170] |
| EfM3.006460 | - | 1.3 | **-8.4** | 1.1 | **-20.3** | **-7.06** | No | KZZ96531 | hypothetical protein AAL_03760 [Moelleriella libera RCEF 2490] |
| EfM3.007440 | - | 1.2 | -1.5 | -1.4 | **-2.3** | -1.32 | No | XP_018140327 | hypothetical protein VFPPC_11200 [Pochonia chlamydosporia 170] |
| EfM3.007580 | - | 1.3 | **3.6** | 1.0 | **3.1** | **15.08** | No | KFG80511 | hypothetical protein MANI_022663 [Metarhizium anisopliae] |
| EfM3.007740 | - | 1.7 | 1.1 | **2.7** | **-3.0** | 2.28 | No | CCE34646 | uncharacterized protein CPUR_08580 [Claviceps purpurea 20.1] |
| EfM3.008680 | - | 1.8 | 1.4 | 1.2 | **2.2** | 1.23 | Yes | NA | |
| EfM3.008740 | - | **-2.8** | 1.7 | -1.1 | **2.7** | **4.04** | Yes | NA | |
| EfM3.009460 | - | 1.3 | 1.2 | -1.0 | -1.1 | **6.46** | No | XP_018140951 | hypothetical protein VFPPC_09225 [Pochonia chlamydosporia 170] |
| EfM3.014350 | sspO | 1.1 | **-2.5** | -1.0 | **-5.1** | **-2.72** | Yes | KJZ74698 | hypothetical protein HIM_05815 [Hirsutella minnesotensis 3608] |
| EfM3.016770 | sspM | 1.6 | **-2.9** | 1.3 | **-29.3** | **-31.24** | Yes | ELA34989 | hypothetical protein CGGC5_5250 [Colletotrichum fructicola Nara gc5] |
| EfM3.026630 | - | **2.3** | **6.2** | 1.1 | **4.5** | **72.39** | Yes | NA | |
| EfM3.027550 | - | **2.3** | **2.4** | **2.0** | **3.7** | 1.75 | No | NA | |
| EfM3.033250 | - | 1.0 | **6.0** | -1.1 | **4.0** | 2.94 | No | PHH60641 | hypothetical protein CDD81_1392 [Ophiocordyceps australis] |
| EfM3.033650 | - | 1.1 | 1.0 | 1.0 | 1.1 | **3.09** | No | NA | |
| EfM3.034400 | - | 1.1 | 1.1 | -1.2 | **4.1** | 5.58 | No | NA | |
| EfM3.038640 | - | -1.2 | -1.2 | 1.0 | **-2.3** | -1.68 | No | XP_007825160 | hypothetical protein MAA_08971 [Metarhizium robertsii ARSEF 23] |
| EfM3.040200 | - | -1.9 | -1.2 | -1.1 | 1.3 | **41.56** | Yes | XP_007825947 | hypothetical protein MAA_09758 [Metarhizium robertsii ARSEF 23] |
| EfM3.041600 | - | 1.2 | 1.8 | -1.1 | **2.2** | **16.36** | No | XP_018139722 | hypothetical protein VFPPC_14197 [Pochonia chlamydosporia 170] |
| EfM3.041760 | - | 1.4 | **2.0** | -1.0 | 1.3 | 2.42 | No | NA | |
| EfM3.041770 | - | 1.1 | 1.8 | 1.0 | **4.7** | **55.32** | Yes | NA | |
| EfM3.042450 | - | -1.7 | **-2.2** | -1.5 | **-2.1** | -1.02 | No | PVI00678 | hypothetical protein DM02DRAFT_614138 [Periconia macrospinosa] |
| EfM3.042700 | - | -1.9 | **3.0** | 2.0 | **2.6** | **2.76** | No | CCE27014 | uncharacterized protein CPUR_00486 [Claviceps purpurea 20.1] |
| EfM3.043290 | - | 1.2 | 1.2 | -1.1 | **2.4** | 1.36 | No | XP_018176067 | hypothetical protein VFPFJ_07599 [Purpureocillium lilacinum] |
| EfM3.045040 | - | 1.1 | **-2.9** | -1.0 | -1.1 | -2.12 | No | NA | |

| | | | | | | | | | |
|---|---|---|---|---|---|---|---|---|---|
| EfM3.048790 | - | **2.6** | 1.8 | 1.5 | 1.5 | -2.07 | No | NA | |
| EfM3.048990 | - | -1.2 | 2.0 | **2.2** | **3.8** | -5.02 | Yes | NA | |
| EfM3.050590 | - | 1.2 | -1.3 | -1.2 | **2.1** | 1.29 | No | XP_013957240 | hypothetical protein TRIVIDRAFT_222299 [Trichoderma virens Gv29-8] |
| EfM3.050840 | - | **-2.3** | **-6.7** | **-5.5** | **-4.3** | -1.71 | No | CEJ95220 | hypothetical protein VHEMI10714 [Torrubiella hemipterigena] |
| EfM3.051300 | - | **2.1** | 1.7 | 1.2 | 1.6 | **6.73** | No | NA | |
| EfM3.055320 | - | 1.0 | 1.5 | 1.0 | 1.8 | **2.48** | No | CCE29234 | related to cytosolic Cu/Zn superoxide dismutase [Claviceps purpurea 20.1] |
| EfM3.057230 | - | -1.6 | **-5.8** | -1.6 | 1.6 | **11.90** | No | XP_023425767 | uncharacterized protein |
| EfM3.058140 | - | 1.2 | **2.9** | 1.1 | **2.6** | 1.94 | No | CZS94271 | uncharacterized protein FFUJ_05161 [Fusarium fujikuroi IMI 58289] |
| EfM3.060210 | - | 1.4 | 1.0 | -1.4 | **2.8** | 5.65 | No | NA | |
| EfM3.061720 | - | -1.0 | 1.8 | -1.4 | 1.0 | **-2.35** | No | NA | |
| EfM3.062700 | sspN | -1.3 | **-56.3** | 1.3 | **-37.4** | **-99.84** | Yes | PHH63509 | hypothetical protein CDD81_5790 [Ophiocordyceps australis] |
| EfM3.062880 | - | 0.0 | **-7.0** | 1.0 | -2.7 | -1.43 | No | CCE29635 | uncharacterized protein CPUR_03482 [Claviceps purpurea 20.1] |
| EfM3.067730 | - | 1.5 | 1.4 | 1.0 | **3.0** | -1.29 | No | NA | |
| EfM3.068330 | - | -1.2 | **-2.8** | **-2.1** | **-3.5** | **-2.52** | No | NA | |
| EfM3.068730 | - | 1.9 | 1.3 | -1.1 | 1.9 | **6.86** | No | NA | |
| EfM3.069580 | - | -1.2 | **-3.2** | -1.7 | **-5.4** | **-2.52** | No | CCE33891 | uncharacterized protein CPUR_07819 [Claviceps purpurea 20.1] |
| EfM3.072390 | - | -1.1 | **2.5** | -1.1 | **5.1** | **4.81** | Yes | NA | |
| EfM3.075190 | - | **3.0** | 1.5 | 1.5 | 2.1 | -2.10 | No | OAA38301 | hypothetical protein NOR_06691 [Metarhizium rileyi RCEF 4871] |
| EfM3.075450 | - | -1.5 | -2.0 | -1.2 | **-2.1** | -1.13 | No | CCE33525 | uncharacterized protein CPUR_07450 [Claviceps purpurea 20.1] |
| EfM3.075920 | - | 1.9 | **12.6** | 1.6 | **3.6** | 7.69 | Yes | CCE31639 | uncharacterized protein CPUR_05492 [Claviceps purpurea 20.1] |
| EfM3.076450 | - | 2.0 | **3.6** | 1.4 | 1.8 | **-2.13** | No | NA | |
| EfM3.077900 | - | **2.6** | **2.2** | 1.6 | 1.8 | **8.22** | No | CCE29593 | uncharacterized protein CPUR_03440 [Claviceps purpurea 20.1] |
| EfM3.079420 | - | -1.0 | **-5.7** | 1.0 | -1.4 | 1.69 | No | XP_018139353 | fungal hydrophobin domain-containing protein [Pochonia chlamydosporia 170] |
