## Supplemental Figures for "VelA and LaeA are key regulators of the *Epichloë festucae* transcriptomic response during symbiosis with perennial ryegrass"

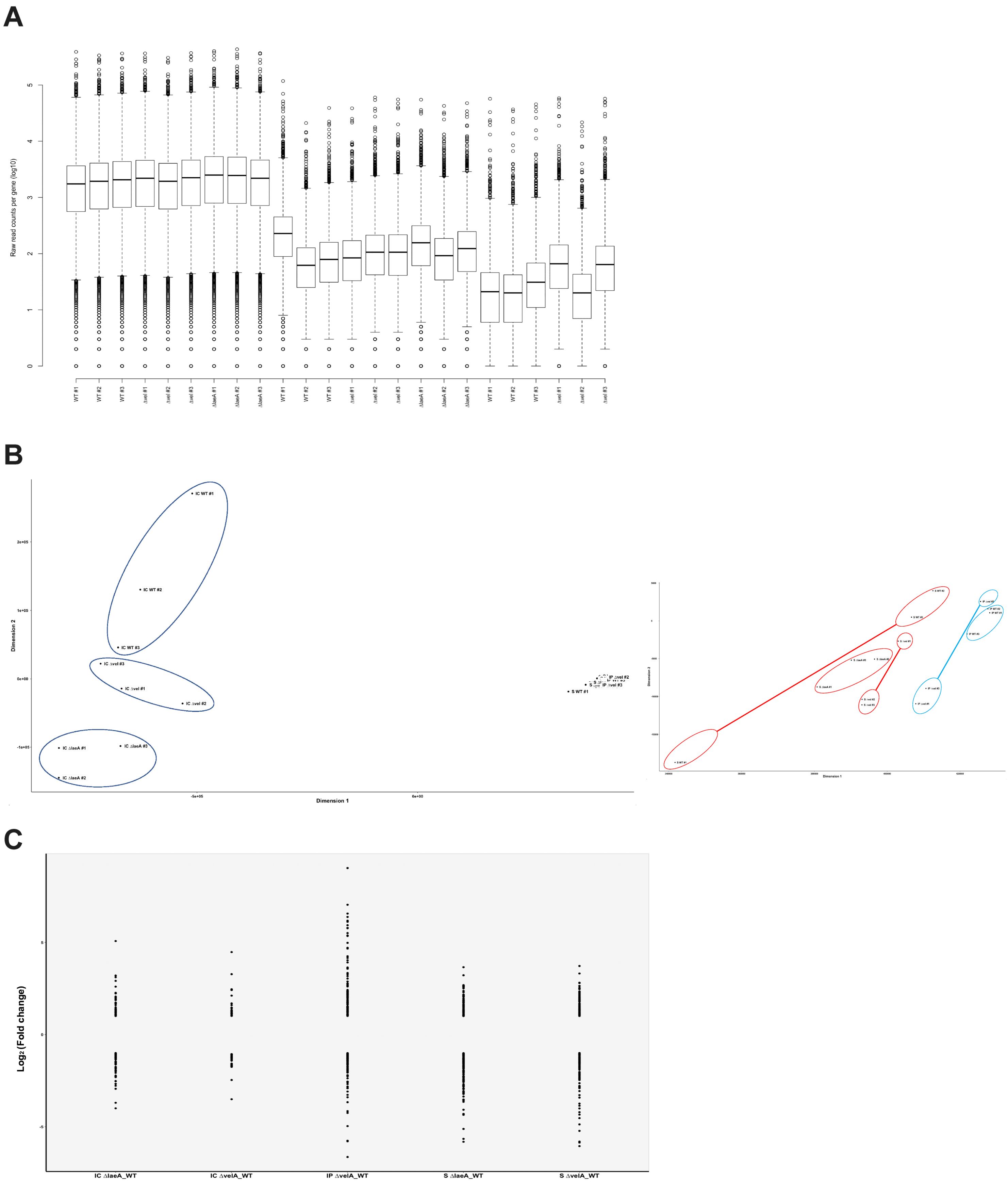


**Fig. S1-** Distribution of number of mapped reads in different samples and fold change in DEGs. A) The bar chart shows the distribution of raw reads mapped per gene. B) Principal component analysis (PCA) of first 1000 highly expressed genes in different samples. C) Fold change distribution of DEGs in different comparisons.

A

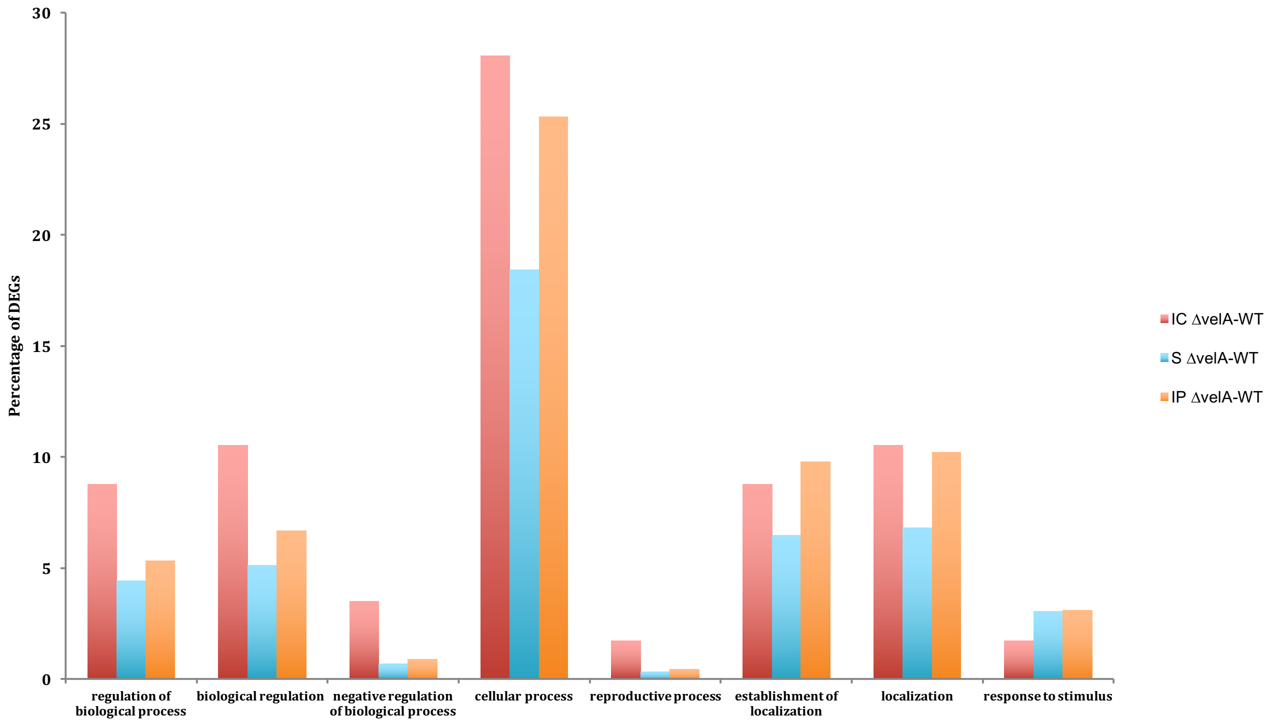


B

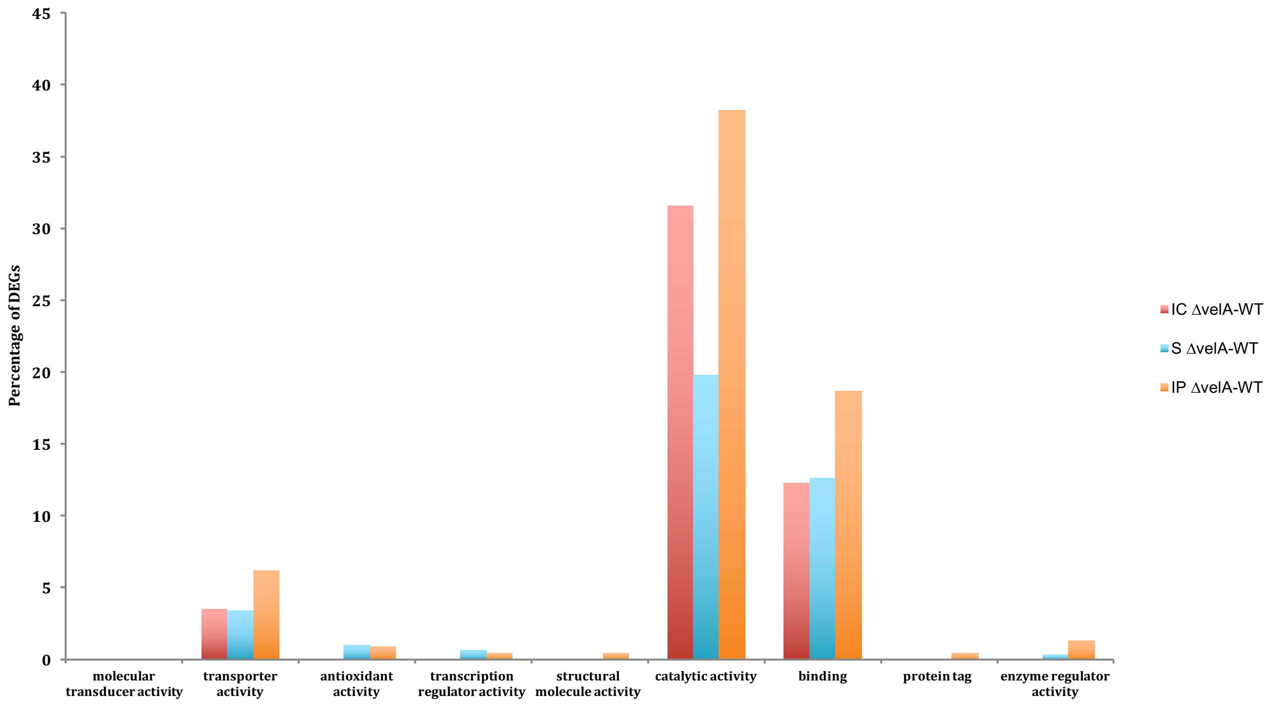


**Fig. S2-** DEGs of different comparisons classified based on primary 'Molecular Function' and 'Biological Process' gene ontology. (a) Bar chart of organised DEGs based on 'Biological Process' GO category. (b) Bar charts of organised DEGs based on 'Molecular Function' GO category. Categories are as follows: GO:0060089 (molecular transducer activity), GO:0005215 (transporter activity), GO:0016209 (antioxidant activity), GO:0030528 (transcription regulator activity), GO:0005198 (structural molecule activity), GO:0003824(catalytic activity), GO:0005488 (binding), GO:0031386 (protein tag), GO:0030234 (enzyme regulator activity).

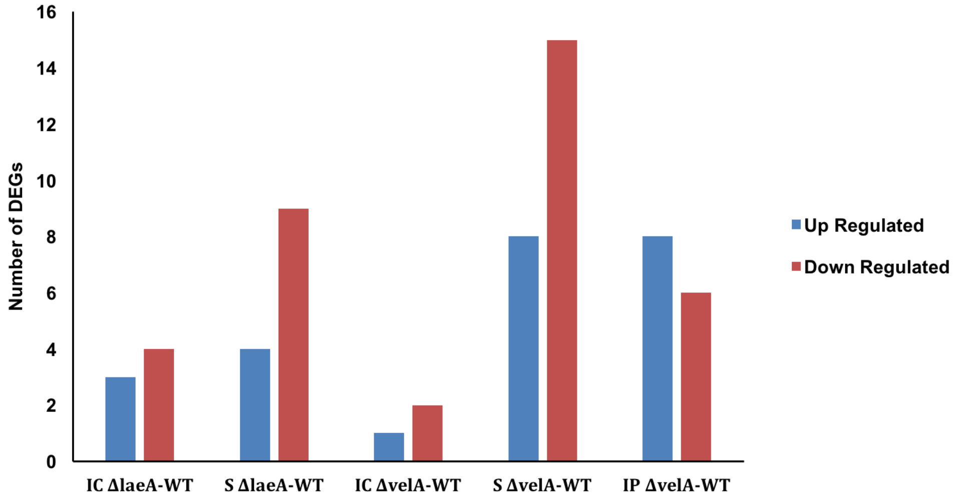


**Fig. S3-** Number of DEGs in different comparisons homologue to CAZyme enzymes in different comparisons.
